# Polarity reversal of stable microtubules during neuronal development

**DOI:** 10.1101/2025.02.07.636940

**Authors:** Malina K. Iwanski, Albert K. Serweta, H. Noor Verwei, Bronte C. Donders, Lukas C. Kapitein

**Affiliations:** Cell Biology, Neurobiology and Biophysics, Department of Biology, Faculty of Science, Utrecht University, Utrecht, The Netherlands

## Abstract

Neurons critically depend on long-distance transport orchestrated by motor proteins walking over their highly asymmetric microtubule cytoskeleton. These microtubules are organized uniformly in axons with their plus-end pointing away from the soma. In contrast, in the dendrites of vertebrate neurons, microtubules are of mixed polarity, but organized into bundles of uniform polarity with stable, long-lived microtubules preferentially oriented minus-end-out and dynamic microtubules oriented plus-end-out. This organization is thought to be essential for guiding selective transport into dendrites, yet how this organization is established is unclear. Here we use a combination of single molecule localization microscopy, expansion microscopy, and live-cell imaging to examine how the microtubule cytoskeleton is reorganized during neuronal development of cultured rat hippocampal neurons. We find that, while the youngest neurites contain microtubules of mixed polarity, stable microtubules are initially preferentially oriented plus-end-out. At this stage of development many stable microtubules are connected to the centrioles, providing an explanation for their plus-end out orientation in emerging neurites. In later stages, these microtubules are released from the centrioles and reorient by sliding between or within neurites to become progressively more minus-end-out. Moreover, prior to axon specification, we commonly observed already one or two minor neurites with an almost uniformly plus-end-out microtubule network, indicative of transient polarization. Together, our findings reveal how stable microtubules are reorganized to help establish the stereotypical microtubule networks seen in the axon and dendrites of mature vertebrate neurons.

## INTRODUCTION

For their proper function, many cells in our body must be able to establish and maintain a polarized organisation. Of these cells, one of the most exquisite examples is neurons, branched cells which bear both structurally and functionally distinct processes known as axons and dendrites. Information is received via the dendrites and integrated, after which an output is generated that is transmitted to other cells via the axon. For this functional distinction, these compartments also require a highly asymmetric distribution of specific molecular players. Thus, which cargo is transported into and out of these processes must be closely regulated. This long-distance cargo transport is effectuated by motor proteins of the kinesin superfamily and dynein that move cargoes unidirectionally, helping to specifically localize them to one or the other compartment. Kinesin and dynein do so by walking along microtubules, intrinsically polar polymers that can grow to many micrometers in length. Microtubules are comprised of α,β-tubulin heterodimers that polymerize head-to-tail to form filaments with a distinct highly dynamic plus-end and a lesser dynamic and often capped minus-end. Most members of the kinesin superfamily walk towards the microtubule plus-end, whereas dynein walks towards the minus-end. In the axons of vertebrate neurons, microtubules are uniformly oriented with their plus-ends outwards away from the soma, whereas dendrites have a mixed microtubule array (Baas et al., 1988). This means that in the axon, most kinesin motors act as anterograde motors, carrying cargo away from the soma, while dynein serves as a retrograde motor. In the dendrites, however, both kinesins and dynein can theoretically act as anterograde and retrograde transporters and additional layers of regulation are required to properly direct these motors. For example, microtubule-associated proteins (MAPs) have distinct activating/inhibitory effects on different motor proteins and some of these are specifically localized to dendrites and may thus help guide this transport (Karasmanis et al., 2018; Monroy et al., 2020). It has also been demonstrated that motors prefer microtubules bearing different tubulin post-translational modification (PTMs), with kinesin-1, for example, preferring microtubules marked by acetylation and detyrosination (typical of long-lived, stable microtubules) and kinesin-3 preferring microtubules marked by tyrosination (typical of labile microtubules with rapid turnover) (Cai et al., 2009; Tas et al., 2017).

Neurons are not directly established with this spectacularly asymmetric array of microtubules properly decorated by MAPs and PTMs. Instead, it is built as the cells polarize during development. The development of dissociated embryonic hippocampal neurons in culture is classically described in five stages (**Figure 1A**) (Dotti et al., 1988). First, once the globular cells in suspension adhere to the coverslip, they extend a lamellipodium, a large protrusion with an extensive branched actin network (stage 1). Then, the cells develop minor processes called neurites that grow and shrink (stage 2). One of these neurites is then specified to become the axon, growing out rapidly and persistently to become much longer than the rest (stage 3). Next, the remaining neurites also branch and develop further to adopt a dendritic identity (stage 4). Finally, the neurons form mature synapses with one another and are fully developed (stage 5). Concomitantly with this process, the microtubule cytoskeleton also adopts a highly polar architecture (**Figure 1A**). In stage 1, the centrosome is still active and the cells have a largely radial microtubule array; however, the centrosome is inactivated later during the developmental process (Stiess et al., 2010). Furthermore, based on the tracking of end binding (EB) protein comets at the growing plus-ends of microtubules, it is known that neurites in stage 2 and 3 neurons contain microtubules of mixed orientation (Yau et al., 2016), but their precise organization remains unclear. In stage 4 and 5 neurons, it is known that axons contain uniformly plus-end-out microtubules, regardless of whether these microtubules are stable or labile (Baas et al., 1988). In the dendrites, however, stable microtubules are preferentially oriented minus-end-out and enriched centrally, while dynamic or labile microtubules are preferentially oriented plus-end-out and enriched peripherally (Katrukha et al., 2021; Tas et al., 2017). This raises the question: how does this organization emerge? Specifically, are microtubules in the minor neurites of stage 2 and 3 neurons already organized into bundles of preferred polarity as is observed in mature dendrites (Tas et al., 2017)? Do stable microtubules in these minor neurites have a preferred minus-end-out orientation as is observed in mature dendrites (Tas et al., 2017) and as has been suggested for developing neurons (Yau et al., 2016)? What are the effects of centrosome inactivation and how do microtubules transition from the radial array to help form the first emerging neurites as neurons transition from stage 1 to stage 2?

**Figure 1:**
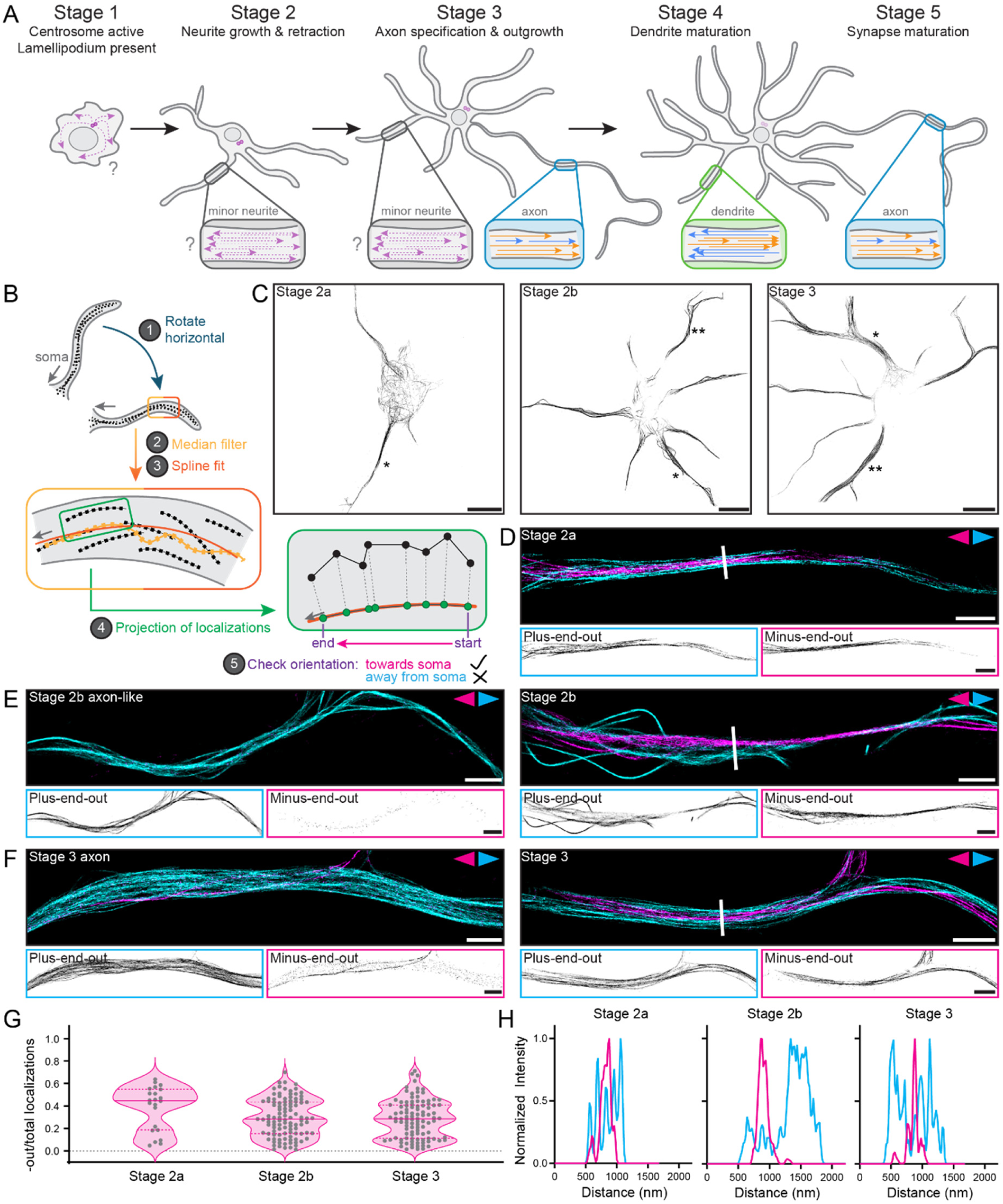
Microtubules of opposite orientation are segregated from early on in neuronal development. (**A)** Schematic showing the stages of neuronal development and the accompanying changes in the microtubule cytoskeleton. While it is known that microtubules are of mixed polarity (indicated by arrowheads) in minor neurites early on in development, the localization and orientation of stable (orange) and labile (blue) microtubules are unclear. **(B)** Schematic showing the step of the analysis pipeline that allows for pseudo-colouring of tracks based on whether they are oriented towards (magenta) or away from (cyan) the soma. **(C)** Total tracks after filtering (all orientations) of representative neurons in early stage 2 (stage 2a), late stage 2 (stage 2b), and stage 3. Scale bars 10 µm. * indicates neurites used in D, E, F right. ** indicates neurites used in E, F left. **(D)** A motor-PAINT reconstruction of a neurite from the stage 2a cell in C with microtubules pseudo-coloured based on whether their plus-end is oriented towards (magenta) or away from (cyan) the soma. Single channel images are shown below. Scale bars 2 µm. **(E)** Motor-PAINT reconstructions of two neurites from the stage 2b cell in C with microtubules pseudo-coloured based on whether their plus-end is oriented towards (magenta) or away from (cyan) the soma. Single channel images are shown below. Scale bars 2 µm. **(F)** Motor-PAINT reconstructions of two neurites from the stage 3 cell in C with microtubules pseudo-coloured based on whether their plus-end is oriented towards (magenta) or away from (cyan) the soma. Single channel images are shown below. Scale bars 2 µm. **(G)** Quantification of the fraction of localizations constituting minus-end-out tracks over the total amount of localizations for neurons in stages 2a, 2b, and 3. Each dot represents one neurite. Medians (0.45, 0.28, 0.29) and interquartile ranges ((0.19, 0.55), (0.15, 0.44), (0.11, 0.41)) are shown. n = 22, 105, 103 neurites from N = 9, 18, 17 cells for stages 2a, 2b, and 3. **(H)** Intensity profiles of minus-end-out (magenta) and plus-end-out (cyan) microtubules along the white lines across the neurites shown in D, E (right), and F (right). Normalization was done independently for the two orientations using the minimum and maximum values of those data sets.

Here, we address these questions using motor-PAINT to analyze the orientation of microtubules at different stages during neuronal development. Motor-PAINT is a single molecule localization technique in which purified kinesin motors are allowed to move across the microtubule cytoskeleton of permeabilized and semi-fixed cells to provide both a super-resolved image of the microtubule network and the orientation of these filaments based on the direction of movement of the kinesin motors (Tas et al., 2017). This technique is particularly suitable because it provides insight into the organization of all microtubules, unlike EB comet tracking (Stepanova et al., 2003), which selectively reveals the orientation of growing microtubules. In addition, it is much higher throughput than the electron microscopy-based hook decoration method (Baas et al., 1988). Our work revealed that, while some neurites in stage 2 neurons contain almost exclusively plus-end-out microtubules, most emergent neurites typically contain both minus-end-out and plus-end-out microtubules, with microtubules of opposite orientation being segregated very early on. Furthermore, expansion microscopy revealed that, much like in mature dendrites, stable and labile microtubules, marked by acetylation and tyrosination respectively, also form segregated networks from early on in development. The orientation of stable microtubules is predominantly minus-end-out from later in stage 2 onwards, but interestingly, it is preferentially plus-end-out in early stage 2. This suggests that stable microtubules are initially nucleated at the centrosome and then released during stage 2, allowing them to reverse their orientation once they have a free minus-end. Consistent with this, expansion microscopy revealed stable microtubules emanating from the centrioles in stage 1 neurons, whereas in stage 2 and 3 neurons, we mostly observed short remnants of stable microtubules near the centrioles. Finally, we performed timelapse imaging using StableMARK, a recently established live-cell marker for stable microtubules (Jansen et al., 2023), and observed sliding and polarity reversal of stable microtubules during stage 2. Together, this work provides insights into the reorganization of the microtubule cytoskeleton during neuronal development and has implications for how the axonal and dendritic identities are established.

## RESULTS

### An optimized motor-PAINT protocol aids reconstruction quality and data interpretation

To robustly analyze microtubule orientation in developing neurons, optimization of the original motor-PAINT protocol (Tas et al., 2017) was necessary (**Figure S1**). Some of these adjustments have also been implemented in our recent motor-PAINT studies using MINFLUX and lattice light-sheet microscopy (Deguchi et al., 2023; Iwanski et al., 2023). First, we noticed that microtubules in young neurons sometimes wobbled during acquisitions, blurring the resulting reconstructions. To combat this, we additionally added a small amount (0.04%) glutaraldehyde during the fixation step. Furthermore, we avoided the transfection of neurons with fluorescently-labelled tubulin (e.g. mCherry-tubulin) as this requires electroporation in young neurons and limits the number of cells that can be imaged. Instead, we incubated neurons with phalloidin to allow us to easily find cells and concomitantly facilitate the identification of their developmental stage. During this step, we also added fluorescently-labelled beads to allow for rapid and efficient drift correction. To improve localization precision and track length, we switched to a SNAP-tagged kinesin motor (DmKHC-SNAP), allowing us to take advantage of bright, photostable dyes such as Janelia Fluor (JF) 646. In addition, this motor was added in bulk rather than locally which helped minimize microtubule wiggling during acquisitions (due to a lower local concentration of motors) and allowed us to seal our imaging chambers. This sealing greatly increased the lifetime of the assays and thus our throughput, likely by minimizing oxygen exposure, which in turn helps limit the acidification of the buffer by the glucose oxidase-catalase oxygen scavenger system that could otherwise reduce buffer quality such that motors are no longer able to walk well.

We also optimized the analysis and visualization of our results to improve reconstruction quality in the dense microtubule arrays of the neurites and simplify the interpretation of the resulting reconstructions (see Materials & Methods). In addition to the robust drift correction facilitated by the inclusion of beads, we also improved the reconstructions by using TrackMate (Tinevez et al., 2017) to localize and track the kinesin motors as it has tracking algorithms (e.g., the overlap tracker) that are more suitable for the dense microtubule networks in neurites. Furthermore, we adjusted our filtering to increase the amount of tracks retained (and thereby the coverage of the microtubule network), most notably by carefully and locally filtering for angles between track “steps” to remove only portions of the tracks with highly divergent angles signifying paused motors or reversals in direction resulting from erroneously linked spots (see Materials & Methods). Additionally, we predominantly colour-coded tracks not by their direction in the field of view, but rather by whether they were oriented towards or away from the soma (**Figure 1B**, see **Figure S2** for examples). This orientation assignment was performed as follows: after selecting each neurite, it was rotated approximately horizontal to reduce the number of instances in which a track contained multiple y-values for a given x-value. To produce a midline for a given neurite/branch, the localizations from all the tracks within this neurite/branch were then treated as a whole, median filtered and spline fit. After determining which end of this midline is closer to the soma, the localizations of each track were projected onto this midline and the start- and end-point of the track were compared to determine whether the track was moving towards or away from the soma along the midline. This was more reliable than directly comparing whether the start- or end- point of the track was closer to the soma, especially in instances where neurites or their branches were curled back towards the soma.

### Microtubules are segregated by orientation from early on in development

With these improvements, we could reliably reconstruct the microtubule networks in neurons throughout their development, focusing on the period approximately 5 to 75 hours after plating (**Figure 1C**, **Figure S2**). These neurons were predominantly in stages 1 to 4, with most neurons in stage 1 or 2 at earlier timepoints and stage 2 to 4 at later timepoints. We chose to focus on neurons in stages 2 and 3, but subdivided those in stage 2 into early (stage 2a) and late (stage 2b) to better describe the changes in microtubule organization occurring early on in development (see Materials & Methods). Most neurites had microtubules of both orientations and showed segregation of microtubules of opposite orientations, including those of stage 2a neurons, which were most commonly found a mere 5 hours after plating (**Figure 1D, E-F (right), G-H**). While we observed a wide spread in the fraction of minus-end-out microtubules per neurite at all developmental stages, neurites in stage 2b and 3 neurons had on average fewer minus-end-out microtubules than those of stage 2a neurons, with median values of 0.45, 0.28, and 0.29 for stage 2a, 2b, and 3 neurons, respectively (**Figure 1G**). This is in agreement with earlier work suggesting that neurites initially contain microtubules of mixed polarity (Yau et al., 2016) and demonstrates that developing neurites have overall less minus-end-out microtubules than the ∼50% reported for mature dendrites (Tas et al., 2017). Furthermore, the observed segregation indicates that the mechanisms that establish spatial segregation by orientation, likely involving MAPs or motors with a preference for crosslinking parallel microtubules, must already be operational early on in development (see Discussion).

### Some late stage 2 neurons have an axon-like neurite with a uniformly plus-end-out microtubule array

In addition, we observed neurites (one or two per cell) with almost exclusively plus-end-out microtubules (**Figure 1E-F (left)**, **Figure S2**). While this is expected for axons (of stage 3 neurons), it was interestingly also the case for some stage 2b neurons, in which we could find neurites with predominantly or exclusively plus-end-out microtubules. These neurites of uniform polarity were not necessarily the longest neurite in stage 2b neurons and this correlation emerged more prominently only later in development (**Figure S3**). Importantly, we cannot determine whether the axon-like neurites in stage 2b neurons actually go on to become the axon or whether this is a transient state, as reported earlier (Burute et al., 2022; Jacobson et al., 2006). Furthermore, the appearance of these axon-like neurites might explain why stage 2b and stage 3 neurons had similar levels of minus-end-out microtubules (median values of 0.28 and 0.29) (**Figure 1G**). As reported previously (Yau et al., 2016), axons of stage 3 neurons still contained a few remaining minus-end-out microtubules. Sometimes these axons and axon-like neurites had a small bundle of minus-end-out microtubules proximal to the soma (**Figure S4**). This suggests that plus-end-out uniformity emerges distally first in these neurites, perhaps by retrograde sliding of these minus-end-out microtubules (see Discussion). In light of these axon-like neurites, we also compared the fraction of minus-end-out microtubules per neurite while excluding axons and axon-like neurites, which resulted in overall similar distributions, but slightly higher median values for later developmental stages (0.30 and 0.33 for stage 2b and 3, respectively) (**Figure S5**).

### Stable and labile microtubules are segregated from early on in development

In the dendrites of mature neurons, there is a relationship between the stability of a microtubule and its orientation, with stable microtubules preferentially oriented minus-end-out (Tas et al., 2017). To determine at which stage this organization emerges, we fixed neurons at similar time points after plating to those described above, stained for tyrosinated tubulin (a marker for labile microtubules) and acetylated tubulin (a marker for stable microtubules), and performed Ultrastructure Expansion Microscopy (U-ExM; (Gambarotto et al., 2019)). This revealed that, much like microtubules of opposite orientation (**Figure 1**), microtubules decorated by different PTMs also segregated into different bundles in stage 2 and 3 (minor) neurites (**Figure 2A-F**, **Figure S6**). To quantify this segregation, we exploited the improved z-resolution of expansion microscopy to construct radial distribution maps of the intensities of these two PTMs and averaged these across (minor) neurites (Katrukha et al., 2021). This revealed that acetylated and tyrosinated microtubules indeed have different radial distributions, with acetylated microtubules being, on average, enriched centrally compared to tyrosinated microtubules, particularly later in development (**Figure 2D, F**). In stage 2a cells, the segregation was present, but less evident in the averaged profile. This is likely because, while we observed the two subsets to be segregated at this stage, we did not always see the central-peripheral distinction in these early neurites (**Figure 2A, B**), similar to what we observed with the segregation of microtubules of opposite orientation (**Figure 1H**). Thus, the bundles of acetylated (stable) microtubules might indeed correspond to the bundles of minus-end-out microtubules, while the bundles of tyrosinated (labile) microtubules might correspond to bundles of plus-end-out microtubules.

**Figure 2:**
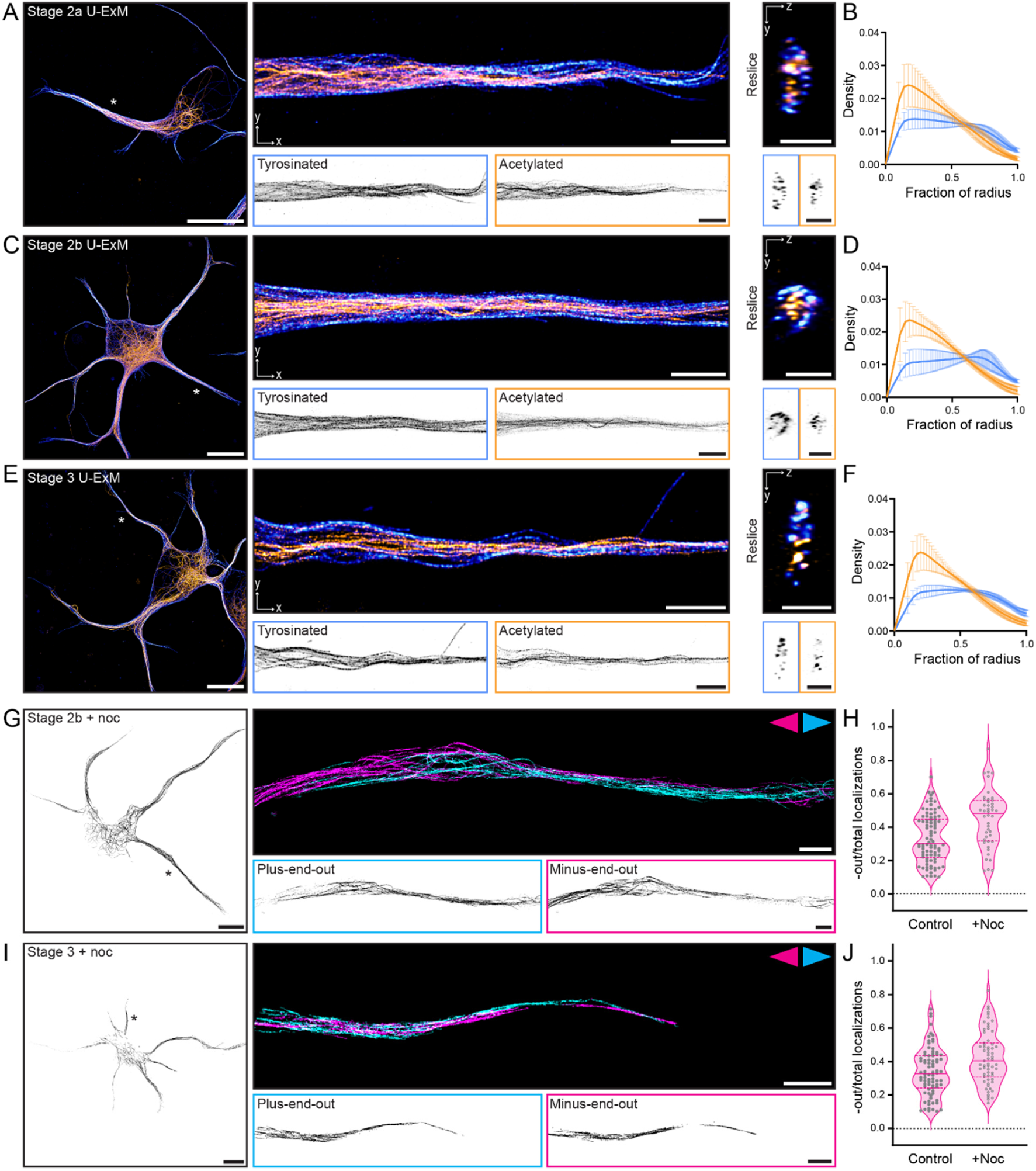
Stable and labile microtubules are segregated from early on in neuronal development, with stable microtubules predominantly oriented minus-end-out in the minor neurites of stage 2b and 3 neurons. **(A)** Left: U-ExM image of a stage 2a cell showing both acetylated (orange) and tyrosinated (blue) microtubules. Scale bar 10 µm (corrected for expansion). * indicates neurite shown to right. Middle: U-ExM image of a single neurite from this cell. Single channel images are shown below. Scale bar 2 µm (corrected for expansion). Right: Y-Z cross-section of the neurite shown. Scale bar 1 µm (corrected for expansion). **(B)** Radial distribution of the intensities of acetylated (orange) and tyrosinated (blue) microtubules averaged across n = 9 neurites from N = 6 neurons. **(C)** Left: U-ExM image of a stage 2b cell showing both acetylated (orange) and tyrosinated (blue) microtubules. * indicates neurite shown to right. Middle: U-ExM image of a single neurite from this cell. Single channel images are shown below. Right: Y-Z cross-section of the neurite shown. Scale bars as in A. **(D)** Radial distribution of the intensities of acetylated (orange) and tyrosinated (blue) microtubules averaged across n = 11 neurites from N = 5 neurons. **(E)** Left: U-ExM image of a stage 3 cell showing both acetylated (magenta) and tyrosinated (cyan) microtubules. * indicates neurite shown to right. Middle: U-ExM image of a single neurite from this cell. Single channel images are shown below. Right: Y-Z cross-section of the neurite shown. Scale bars as in A. **(F)** Radial distribution of the intensities of acetylated (orange) and tyrosinated (blue) microtubules averaged across n = 8 neurites from N = 5 neurons. **(G)** Left: Total tracks after filtering (all orientations) of a stage 2b neuron after nocodazole treatment. Scale bar 10 µm. * indicates neurite shown to right. Right: A motor-PAINT reconstruction of a neurite from this cell with microtubules pseudo-coloured based on whether their plus-end is oriented towards (magenta) or away from (cyan) the soma. Single channel images are shown below. Scale bars 2 µm. **(H)** Quantification of the fraction of localizations constituting minus-end-out tracks over the total amount of localizations for non-axon-like neurites in stage 2b neurons without (control) and with (+ noc) a nocodazole treatment. Each dot represents one neurite. Medians (0.30, 0.48) and interquartile ranges ((0.22, 0.49), (0.32, 0.56)) are shown. n = 90, 44 neurites from N = 18, 11 cells for control and nocodazole-treated cells. **(I)** Left: Total tracks after filtering (all orientations) of a stage 3 neuron after nocodazole treatment. Scale bar 10 µm. * indicates neurite shown to right. Right: A motor-PAINT reconstruction of a neurite from this cell with microtubules pseudo-coloured based on whether their plus-end is oriented towards (magenta) or away from (cyan) the soma. Single channel images are shown below. Scale bars 2 µm. **(J)** Quantification of the fraction of localizations constituting minus-end-out tracks over the total amount of localizations for non-axon neurites in stage 3 neurons without (control) and with (+ noc) a nocodazole treatment. Each dot represents one neurite. Medians (0.33, 0.40) and interquartile ranges ((0.24, 0.44), (0.31, 0.56)) are shown. n = 82, 63 neurites from N = 17, 14 cells for control and nocodazole-treated cells.

To directly query the orientation of stable microtubules, we selectively removed labile microtubules using nocodazole before performing motor-PAINT (Tas et al., 2017). We first verified the efficacy and non-lethality of the nocodazole treatment on neurons early on in development and saw that we could indeed selectively depolymerize the tyrosinated (labile) microtubules, while retaining the acetylated (stable) microtubules (**Figure S7**). Using motor-PAINT, we observed a higher fraction of minus-end-out microtubules in non-axon(-like) neurites after nocodazole treatment as compared to control in stage 2b (median values of 0.30 and 0.48 without and with nocodazole, respectively) and 3 (median values of 0.33 and 0.40 without and with nocodazole, respectively) neurons (**Figure 2G-J**, **Figure S5**, **Figure S8**). We also see similar values for the fraction of stable microtubules in non-axon neurites of stage 4 neurons after nocodazole treatment (medians of 0.32 and 0.45 without and with nocodazole, respectively) (**Figure S9**).This indicates that the stable microtubules in these neurites are indeed preferentially oriented minus-end-out compared to the total population of microtubules, although there are still more plus-end-out stable microtubules during these stages.

### Stable microtubules are oriented plus-end-out in early stage 2 neurites and often connected to the centrosome

Next, we repeated this procedure in stage 2a neurons. It was apparent that there were fewer stable microtubules in these neurons and we observed shorter tracks after nocodazole treatment (**Figure 3A**, **Figure S8**), which could imply that the microtubules are stabilized in shorter stretches at this stage. Interestingly, unlike for stage 2b and 3 neurons, we observed a lower fraction of minus-end-out microtubules in stage 2a neurons after nocodazole treatment as compared to control (median values of 0.45 and 0.18 without and with nocodazole, respectively) (**Figure 3B**). Moreover, four of the six neurites with ≥ 50% minus-end-out microtubules after nocodazole treatment belong to the same cell, making this cell somewhat of an outlier. These results suggest that stable microtubules are initially predominantly oriented plus-end-out. Perhaps stable microtubules are originally nucleated at the centrosome (or microtubules nucleated at the centrosome are stabilized while anchored) such that they are plus-end-out and then later released, for example via severing, to allow them to reverse their orientation to minus-end-out (e.g., via sliding) in dendrites. Indeed, this has been proposed previously for microtubules in both axons and dendrites (Ahmad et al., 1999; Ahmad & Baas, 1995; Baas & Yu, 1996; Sharp et al., 1995) and it is known that the centrosome is initially active in stage 1 neurons and then inactivated during neuronal development preceding axon outgrowth (i.e., stage 3) (Stiess et al., 2010). Alternatively, the plus-end-out stable microtubules could be selectively depolymerized to make way for newly nucleated minus-end-out microtubules, or they could be simply outnumbered by these minus-end-out microtubules.

**Figure 3:**
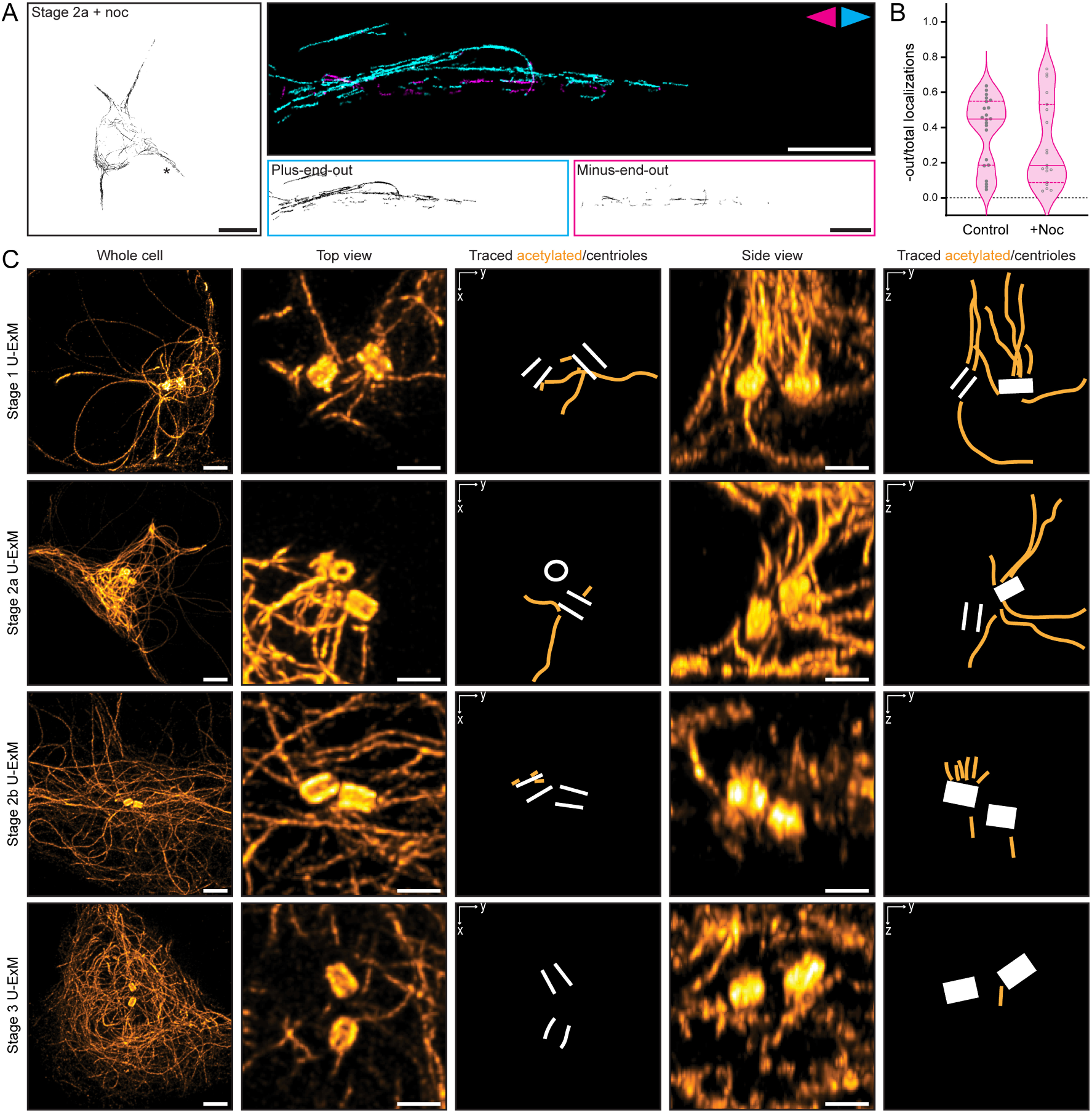
Stable microtubules are predominantly oriented plus-end-out and centrosomally anchored in the emergent neurites of stage 2a neurons. **(A)** Left: Total tracks after filtering (all orientations) of a stage 2a neuron after nocodazole treatment. Scale bar 10 µm. * indicates neurite shown to right. Right: A motor-PAINT reconstruction of a neurite from this cell with microtubules pseudo-coloured based on whether their plus-end is oriented towards (magenta) or away from (cyan) the soma. Single channel images are shown below. Scale bars 2 µm. **(B)** Quantification of the fraction of localizations constituting minus-end-out tracks over the total amount of localizations for stage 2a neurons without (control) and with (+ noc) a nocodazole treatment. Each dot represents one neurite. Medians (0.45, 0.18) and interquartile ranges ((0.19, 0.55), (0.09, 0.53)) are shown. n = 22, 19 neurites from N = 9, 8 cells for control and nocodazole-treated cells. **(C)** Max projections of acetylated microtubules in stage 1, stage 2a, stage 2b, and stage 3 neurons (from top to bottom) imaged using ultrastructure expansion microscopy (U-ExM). Larger overviews of the soma are shown on the left. Scale bars, 1 µm (corrected for expansion). Max projections showing different (X-Y and Y-Z) views of acetylated microtubules around the centrioles are shown in the subsequent columns. Scale bars, 0.5 µm (corrected for expansion). Representations in which the acetylated microtubules directly connected to the centrosome have been traced are also shown.

To study whether stable microtubules are initially anchored at the centrioles, we fixed neurons at similar time points after plating, stained for acetylated tubulin, and again used U-ExM (Gambarotto et al., 2019) to study the microtubule network around the centrioles. Expansion microscopy is well-suited for this because it affords improved z-resolution, which is essential to resolve microtubules in the dense network surrounding the centrioles. This revealed that, early on in development (stage 1 and 2a), long acetylated microtubules were emanating from the centrioles (**Figure 3C**). Later in development, the amount of long acetylated microtubules apparently anchored at the centrioles decreased until predominantly short stumps of acetylated microtubules (< 0.5 µm when corrected for expansion factor) were observed around the centrioles in stage 2b and 3 neurons (**Figure 3C**). Quantifying the absolute length distribution of the anchored microtubules is challenging, but we observed this phenomenon in at least 6 cells from 3 rats for each stage. It was also apparent that, while in stage 1 the microtubules are anchored at both the mother and the daughter centriole, from stage 2a onwards, the microtubules or their remnants were preferentially anchored at one of the two centrioles. The stumps observed around centrioles in stage 2b and 3 neurons could be remnants of severed/released microtubules, suggesting that stable microtubules may be reoriented by sliding after release from the centrosome rather than by depolymerization and repolymerization.

### Stable microtubules undergo extensive sliding and polarity reversal during neuronal development

To better examine how stable microtubules are reorganized during the transition from stage 2a to 2b, we electroporated freshly dissociated neurons with stable microtubule-associated rigor kinesin (StableMARK) (Jansen et al., 2023), a mutant non-motile kinesin-1 motor that specifically binds to stable microtubules, before plating and then imaged between 4 and 27 hours after plating using total internal reflection fluorescence (TIRF) microscopy (**Figure 4A-B**). We observed extensive retrograde flow (**Figure 4C-E**, **Movie 1**) in stage 2b neurons, as has also been observed for the total population of microtubules (Burute et al., 2022; Schelski & Bradke, 2022). The median speed of this flow was 0.15 µm/min (inter-quartile range of 0.10-0.24 µm/min) (**Figure 4E**), in agreement with previously reported values (Burute et al., 2022; Schelski & Bradke, 2022).

**Figure 4:**
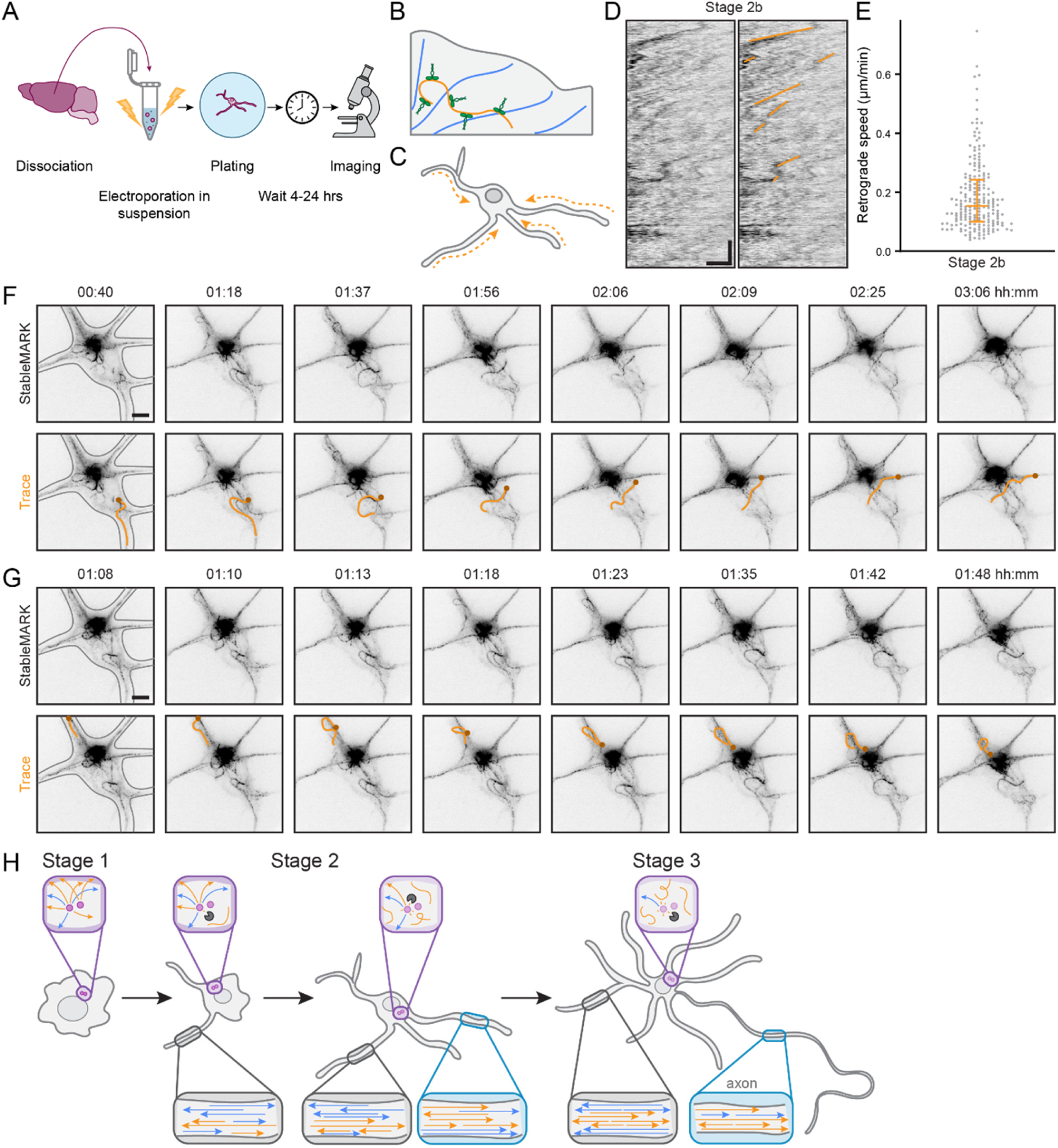
Stable microtubules undergo retrograde flow and reverse their orientation during the early stages of neuronal development. **(A)** Schematic showing how dissociated neurons from embryonic rat hippocampi were electroporated with StableMARK prior to plating on laminin-coated coverslips and imaged shortly thereafter. **(B)** Schematic showing StableMARK (green) bound to stable microtubules (orange), while labile microtubules (blue) are undecorated. **(C)** Schematic showing the retrograde flow of stable microtubules observed in young neurons. **(D)** Kymographs of this retrograde flow in a stage 2b neuron (left) with examples of the lines used to determine the speed of this retrograde flow in orange (right). Scale bars 2 µm (horizontal) and 30 min (vertical). Corresponds to Movie 1. **(E)** Quantification of the speed of retrograde flow from n = 30 neurites in N = 7 stage 2b neurons. Each dot represents one line as traced in D. The median (0.15 µm/min) and interquartile range (0.10 µm/min, 0.24 µm/min) are shown. **(F)** An example of a stable microtubule in a stage 2b neuron reversing its orientation by exiting from one neurite and entering another. The intended microtubule is traced in magenta in the bottom row, with the same end marked by a small circle in each frame to indicate orientation. First frame also has the cell outline in grey. Times shown above each frame are in hours:minutes. Scale bar 5 µm. Corresponds to Movie 5. **(G)** An example of another stable microtubule in the same neuron reversing its orientation by turning back on itself within one neurite. The intended microtubule is traced in orange in the bottom row, with the same end marked by a small circle in each frame to indicate orientation. First frame also has the cell outline in grey. Times shown above each frame are in hours:minutes. Scale bar 5 µm. Corresponds to Movie 5. **(H)** Stable microtubules reverse their orientation during neuronal development by dissociating from the centrosome. Schematic showing the stages of neuronal development and the accompanying changes in the microtubule cytoskeleton. It is now apparent that microtubules of opposite orientation (indicated by arrowheads) are segregated from early on in development, but that stable microtubules (orange) are initially mostly plus-end-out in stage 2a and later reverse their orientation to become minus-end-out. This is facilitated by their detachment from the centrosome (purple), which allows them to be slid into neurites with their minus-ends leading. Labile microtubules (blue) remain largely plus-end-out throughout development.

In addition, we observed highly motile stable microtubules that exhibited extensive sliding in the soma and neurites, particularly in stage 2a and 2b neurons (**Figure S10**, **Movies 2, 3**). There were also instances in which a bundle of stable microtubules from one neurite appeared to be directly linked to a focal point (likely the centrosome) and instances in which the ends of some of these microtubules appeared to detach from this point (**Figure S10**, **Movies 3, 4**), in agreement with our expansion data (**Figure 3B-D**). Importantly, we observed many instances of microtubule polarity reversal, particularly in stage 2b neurons. For example, we could observe a stable microtubule exiting from one neurite and being slid towards another neurite, which it then entered with the opposite orientation (**Figure 4F**, **Movie 5**), and we could observe a stable microtubule curling back on itself within a given neurite, which also reversed its orientation (**Figure 4G**, **Movies 3, 5**). These observations thus demonstrate two means by which stable microtubules can reverse their orientation in late stage 2 neurons. Importantly, we could follow the same microtubule for several hours, arguing against the depolymerization and repolymerization of stable microtubules to reverse their orientation.

## Discussion

In this work, we mapped out the reorganization of the microtubule cytoskeleton that occurs during neuronal polarization (**Figure 4H**). Emerging neurites of early stage 2 neurons already contain microtubules of both orientations and these are typically segregated. These emerging neurites also contain segregated networks of acetylated (stable) and tyrosinated (labile) microtubules. In later stage 2, stage 3, and stage 4 neurons, stable (nocodazole-resistant) microtubules are oriented more minus-end-out compared to the total (untreated) population of microtubules; however, in early stage 2 neurons, stable microtubules are preferentially oriented plus-end-out, likely because their minus-ends are still anchored at the centrioles at this stage. The fraction of anchored stable microtubules decreases during development, while the appearance of short stumps of microtubules attached to the centrioles suggests that these microtubules may be released by severing. Finally, using live-cell imaging we observed the retrograde flow of stable microtubules and two mechanisms by which stable microtubules reverse their orientation. In addition to these phenomena, which predominantly describe the development of the dendrites, we also identified axon-like neurites with (almost) uniformly plus-end-out microtubules in late stage 2 neurons, prior to axon specification.

Our results for microtubule orientations in developing neurites are consistent with earlier work that used EB comet tracking after laser severing in stage 2 and 3 neurons. This work revealed that the fraction of comets moving towards the soma increased after laser severing, which indirectly suggested that minus-end-out microtubules are often more stable (Yau et al., 2016). Our direct analysis of the orientation of stable microtubules revealed very similar fractions of stable minus-end-out microtubules in stage 2b and 3 neurons, but these values are lower than what has been reported for mature neurons (Tas et al., 2017). This suggests that the population of stable minus-end-out microtubules continues to increase during development.

Previous work reported microtubules emanating from the centrioles of developing neurons (Stiess et al., 2010; Yu et al., 1993), but could not selectively detect stable microtubules. Our work now shows that in stage 1 and somewhat in early stage 2, acetylated (stable) microtubules are also anchored at the centrioles. This explains why stable microtubules are predominantly oriented plus-end-out in these stages. During stage 2, many stable microtubules begin to reverse their orientation and are no longer anchored at the centrioles. Instead, we found an increasing density of short acetylated microtubules attached to the centrioles during this stage. Together with the very long lifetime of stable microtubules observed with live-cell imaging, this suggests that stable microtubules are released from the centrosome via severing by spastin, katanin, or fidgetin, followed by redistribution and reorientation. We favor severing over the recently described CAMSAP-mediated release of microtubules from γTuRC (Rai et al., 2024) because of the presence of these microtubule stumps. It is unlikely that CAMSAP, a microtubule minus-end-binding protein, would leave such a microtubule remnant at the centrioles. Taken together, our results show that the initial population of stable microtubules follows the centrosomal-release-and-reorientation model proposed by Baas and colleagues (Baas & Yu, 1996; Yu et al., 1993). In the future, it will be interesting to investigate what triggers the activity of these severing enzymes, as well as the potential effects on microtubule polarity during development in cells depleted of them.

Once released, the plus-end-out stable microtubules must reverse their orientation, whereas the dynamic microtubules do not. We provide evidence that stable microtubules slide between or within neurites to reverse their orientation rather than depolymerize and repolymerize. What remains unclear, however, is how this sliding is effectuated. In the example shown, the microtubule end covers a distance of ∼15 µm in about 4 hours (∼60 nm/min on average). During this time, the microtubule end undergoes bursts of transient motility and periods in which it is rather immobile. We speculate that this motility is driven by microtubule-based motors, perhaps in combination with hitchhiking on a growing plus-end of a dynamic microtubule (Mattie et al., 2010). Multiple motor proteins, which move at a range of speeds, have been implicated in microtubule polarity sorting and sliding during neuronal development (del Castillo et al., 2015; He et al., 2020; Kahn et al., 2014; Lin et al., 2012; Liu et al., 2010; Lu et al., 2013; Mattie et al., 2010; Muralidharan & Baas, 2019; Schelski & Bradke, 2022; Weiner et al., 2020). One of these is kinesin-1, a motor that preferentially binds to and moves on stable microtubules (Cai et al., 2009; Reed et al., 2006) and that has been shown to slide microtubules in multiple cell types including neurons (He et al., 2020; Jolly et al., 2010; Lu et al., 2013; Winding et al., 2016). This motor could crosslink microtubules or crosslink a microtubule with, for example, the actin cortex, and, by walking on the stable microtubule towards its plus-end, push this microtubule with its minus-end leading into a new neurite (Kapitein & Hoogenraad, 2015). Alternatively, a motor that moves along labile microtubules, but that associates with CAMSAPs at the minus-end of a stable microtubule could help drag this minus-end out into a minor neurite. There is indeed evidence that at least the minus-end-directed kinesin-14 can interact with CAMSAP2 (Cao et al., 2020). It is likely that a combination of motors orchestrates the sliding of stable microtubules between neurites and within neurites to facilitate their microtubule polarity reversal. To solve this puzzle, it may be helpful to image StableMARK during neuronal development upon treatment with inhibitors or activators of different motor proteins such as kinesore (activator of kinesin-1) (Randall et al., 2017), monastrol (inhibitor of kinesin-5) (Mayer et al., 1999), or dynarrestin/ciliobrevin D (inhibitors of dynein) (Firestone et al., 2012; Höing et al., 2018).

Within neurites, we observed a clear spatial separation both between labile and stable microtubules and between plus-end-out and minus-end-out microtubules. To achieve this, it is likely important to crosslink microtubules of the same polarity. Interestingly, the microtubule bundler TRIM46 is one of the two MAPs known so far to crosslink microtubules with a preferred relative orientation. It preferentially crosslinks parallel microtubules (Harterink et al., 2019; Van Beuningen et al., 2015), but oscillates between neurites in stage 2 before settling in the axon (Burute et al., 2022), making it an unlikely candidate for bundling parallel microtubules in the minor neurites and dendrites; however, there are other microtubule-binding TRIM family members that may fulfill this role (Glover et al., 2024; Short & Cox, 2006). Parallel bundling by TRIM46 has also been shown to induce stabilization of the bundled microtubules (Van Beuningen et al., 2015). Such bundling-induced stabilization may contribute to the amplification of the initial population of centriole-derived and polarity-reversed stable microtubules, as the number of stable microtubules continues to increase after the inactivation of the centrosome.

Finally, we observed the presence of axon-like neurites with (almost) uniformly plus-end-out microtubule arrays in late stage 2 neurons prior to axon specification. Importantly, these neurites were not always the longest of the cell and from the present study, we cannot deduce whether these will indeed become the axon, but these observations are consistent with earlier reports of transient polarization in this stage (Burute et al., 2022; Jacobson et al., 2006). It will thus be interesting to see whether this plus-end-out uniformity coincides with the waving of kinesin-1 and TRIM46, two axonal markers, that occurs in stage 2 neurons (Burute et al., 2022).

The present study was conducted in dissociated rat hippocampal neurons, an important model system used by many labs worldwide. Nevertheless, it is interesting to also consider how this process may be similar or different *in situ*. In the brain, there are many factors involved in directing neuronal migration, differentiation, and axon specification. For example, positive and negative guidance cues help direct axon growth and stiffness gradients are also known to play a role both in neuronal differentiation (Engler et al., 2006) and axon specification (Burute et al., 2022). As such, we expect there to be less variability between neurites and perhaps a more efficient axon specification step such that stage 2 is rather brief. However, given that neurons arise by cell division from neuronal precursor cells, we expect their centrosomes to start as active microtubule nucleating centers and progressively inactivate, much like the dissociated hippocampal neurons we used. This means it is not implausible that stable microtubules are similarly originally nucleated from and/or anchored at the centrosome, making them plus-end-out, and then later released to allow them to reverse their orientation. However, more tools are needed to better examine the organization and orientation of microtubules in tissue to validate this. At present, this study provides the most in-depth examination of microtubule reorganization during development to-date and exploits the relatively easy manipulation and high-resolution imaging of cultured rat hippocampal neurons to gain insights into how stable microtubules are made and re-oriented during the course of development.

## MATERIALS & METHODS

### CELL CULTURE

For neuron cultures, 18 mm coverslips (No. 1.5 high precision) were cleaned and coated with poly-L-lysine (37.5 µg/mL) and laminin (1.25 µg/mL). Subsequently, hippocampi were dissected from embryonic day 18 rat brains (sex not determined) and dissociated. For all experiments without electroporation, the resulting primary hippocampal neurons were then plated at a density of 50, 000 cells/well (12-well plate) onto the treated coverslips in neurobasal medium (NB) supplemented with 2% B27, 0.5 mM glutamine, 15.6 µM glutamate, and 1% penicillin/streptomycin (full medium). These cells were maintained at 37 °C and 5% CO_2_ until use. For more details, see (Kapitein et al., 2010).

### DRUG TREATMENT

Nocodazole treatment was performed as described previously (Tas et al., 2017). Cells were treated with 4 µM nocodazole for 2.5 hours at 37 °C and 5% CO_2_. For immunofluorescence assays, control samples were treated with an equivalent amount of DMSO (∼0.04%). For motor-PAINT assays, no DMSO was added to control samples.

### IMMUNOFLUORESCENCE

To verify the efficiency of our nocodazole treatment, neurons were fixed on DIV0, DIV1, DIV2, and DIV3 as follows. First, soluble tubulin was extracted by subjecting cells to an extraction step (60 s, buffer pre-warmed to 37 °C) with 0.3% Triton X-100 and 0.1% glutaraldehyde in MRB80 (80 mM K-Pipes, 1 mM EGTA, 4 mM MgCl_2_; pH 6.80). This buffer was then exchanged to allow for fixation (10 min, buffer pre-warmed to 37 °C) using 4% formaldehyde and 4% w/v sucrose in MRB80. Cells were then rinsed with PBS, permeabilized using 0.2% Triton X-100 in PBS for 10 min, rinsed with PBS, quenched three times for 5 min in 5 mg/mL sodium borohydride in PBS, rinsed with PBS, and blocked for 30 min in 3% w/v BSA in PBS. Samples were then incubated in primary antibodies (see antibody table) for 1.5-2 hours at room temperature, washed with PBS, and then incubated with secondary antibodies (see antibody table) for 1-1.5 hours. Finally, samples were washed with PBS and then MilliQ water, dried, and lastly mounted in ProLong Diamond Antifade Mountant.

The following antibodies were used in this study:

| **Antibody** | **Company** | **Catalog No.** | **Dilution** | **Use** |
| --- | --- | --- | --- | --- |
| Rat anti-tyrosinated tubulin | Abcam | ab6160 | 1:500<br>1:250 | Confocal<br>U-ExM |
| Rabbit anti-acetylated tubulin | Cell Signaling Technology | 5335 | 1:500<br>1:250 | Confocal<br>U-ExM |
| Alexa Fluor 647 Goat anti-rat | Life Technologies | A-21247 | 1:500 | Confocal |
| Alexa Fluor 488 Goat anti-rabbit | Thermo Fisher Scientific | A-11008 | 1:500<br>1:500 | Confocal<br>U-ExM |
| Alexa Fluor 594 Goat anti-rat | Thermo Fisher Scientific | A-11007 | 1:500 | U-ExM |

### CONFOCAL IMAGING

Fixed samples were imaged on an LSM 700 laser scanning confocal microscope (Zeiss): an Axio Observer Z1 inverted microscope with a motorized stage, a plan-apochromat 63x oil immersion objective (NA 1.40), and 488/555/639 nm laser lines. The following filters were used: bandpass (BP) 490-555 nm, BP 592-622 nm, and long pass (LP) 640 nm. Detection was performed using a multialkali PMT. Components were controlled using ZEN 2012 (Zeiss).

### ELECTROPORATION

Electroporation was performed immediately after dissociation, prior to plating. Dissociated neurons were pelleted (200 ×g, 5 min) and then gently resuspended in Nucleofector^TM^ solution (Amaxa Biosystems/Lonza Bioscience) freshly supplemented with 20% fetal calf serum. The neurons were then mixed with 0.5 µg StableMARK (Jansen et al., 2023; addgene no. 174649) in an electroporation cuvette with 200,000-1,000,000 neurons per reaction. These neurons were then electroporated using the Lonza Nucleofector 2b on the O-003 (rat hippocampal neurons) setting. After adding full medium, these cells were plated on cleaned and coated 25 mm coverslips (No. 1.5 high precision; see above) at a density of 167,000 cells/well (6-well plate).

### LIVE-CELL TIRF IMAGING

For live-cell imaging, neurons on 25 mm coverslips were mounted in imaging rings with conditioned medium and sealed using an additional coverslip to maintain CO_2_ levels. Images were acquired on an inverted Nikon Eclipse Ti-E microscope equipped with a 100 x Apo TIRF oil immersion objective (NA 1.49), a Perfect Focus System, an ASI motorized stage MS-2000-XY, and an iLas2 system (Gataca Systems) for azimuthal TIRF/oblique illumination imaging. For imaging, a Stradus 488 nm (150 mW; Vortran) laser was used together with the ET-GFP filter set (49002; Chroma). Images were projected onto the chip of an Evolve Delta 512 EMCCD camera (Photometrics) with a 2.5 × intermediate lens (Nikon C mount adapter 2.5 ×) at a magnification of 0.065 µm/pixel and 16-bit pixel depth. To keep cells at 37 °C, we used a stage top incubator (model STXG-PLAMX-SETZ21L; Tokai Hit). Images were acquired with 6.5% laser power and 200 ms exposure time with a 1 min interval. Components were controlled using MetaMorph 7.10 (Molecular Devices).

### PROTEIN PURIFICATION

For motor-PAINT, DmKHC[1-421]-SNAP-6xHis (addgene no. 196975) was purified from E. coli BL21 cells as previously described (Deguchi et al., 2023). Briefly, after transformation, bacteria were cultured until OD_600_ ≈ 0.7 at 37 °C and 180 rpm. Cultures were then cooled to < 20 °C, after which protein expression was induced with 0.15 mM IPTG at 18 °C and 180 rpm overnight. Cells were then pelleted by centrifugation at 4500 ×g, snap frozen in liquid nitrogen, and stored at −80 °C until use. On the day of purification, cells were rapidly thawed at 37 °C before being resuspended in chilled lysis buffer (50 mM sodium phosphate buffer supplemented with 5 mM MgCl_2_, 5 mM imidazole, 10% v/v glycerol, 300 mM NaCl, 0.5 mM ATP, and 1 × EDTA-free cOmplete protease inhibitor; pH 8.0). Bacteria were lysed by sonication (5 rounds of 30 s), supplemented with 2 mg/mL lysozyme, and then incubated on ice for 45 min. The lysate was clarified by centrifuging at 26000 ×g for 30 min before being incubated with equilibrated cOmplete His-tag purification resin for 2 hrs. Beads were then pelleted and resuspended in 5 column volumes (CV) wash buffer (50 mM sodium phosphate buffer supplemented with 5 mM MgCl_2_, 5 mM imidazole, 10% v/v glycerol, 300 mM NaCl, and 0.5 mM ATP; pH 8.0) four times. Finally, the resin was transferred to a BioRad column. Once settled, the wash buffer was allowed to elute before adding 3 CV elution buffer (50 mM sodium phosphate buffer supplemented with 5 mM MgCl_2_, 300 mM imidazole, 10% glycerol, 300 mM NaCl, and 0.5 mM ATP; pH 8.0) to elute the protein. The eluent was collected, concentrated by spinning through a 3000 Da MWCO filter, and supplemented with 1 mM DTT and 10% w/v sucrose. These aliquots were then flash frozen in liquid nitrogen and stored at −80 °C. For labelling, protein was thawed and incubated with an additional 1 mM DTT for 30 min before adding 50 μM JF646-SNAP-tag ligand and incubating with rotation overnight. Finally, protein was exchanged into wash buffer (low imidazole) supplemented with 2 mM DTT and 10% w/v sucrose by spinning through a 3000 Da MWCO filter. This also removes excess dye molecules as these are not retained by the filter. Concentration was determined with a BSA standard gel. All steps from lysis onwards were performed at 4 °C.

### MOTOR-PAINT SAMPLE PREPARATION

For this work, the motor-PAINT protocol described previously (F. Chen et al., 2022; Deguchi et al., 2023; Tas et al., 2017) was somewhat modified. Neurons were permeabilized for 1 minute in extraction buffer (BRB80 (80 mM K-PIPES, 1 mM MgCl_2_, 1 mM EGTA; pH 6.80) supplemented with 1 M sucrose and 0.15% v/v TritonX-100) pre-warmed to 37 °C. An equal volume of pre-warmed fixation buffer (BRB80 supplemented with 2% w/v paraformaldehyde (PFA) and 0.08% glutaraldehyde (GA) was added to this (i.e. final PFA concentration of 1% w/v and final GA concentration of 0.04%) and the solutions were mixed by gently pipetting for 1 min. Subsequently, the extraction/fixation buffer mixture was removed, and the sample was washed four times with pre-warmed wash buffer (BRB80 supplemented with 1 μM Taxol) for 1 min. The sample was then incubated with wash buffer with added phalloidin 405 (165 nM or 1:400; Thermo Fisher Scientific A12379) and 0.1 µm TetraSpeck beads (10^8^ beads or 1:1000; Life Technologies T7279; sonicated before use) and incubated at 37 °C for 8-10 min. Next, this solution was removed, and the chamber was washed three times for 1 min with wash buffer. Before adding, an aliquot of DmKHC[1-421]-SNAP motors was thawed and spun in an Airfuge (Beckman Coulter) at 20 psi for 5 min in a pre-chilled rotor to remove any aggregates, transferred to a clean tube, and kept on ice until use. Finally, the wash buffer was exchanged for pre-warmed imaging buffer (BRB80 supplemented with 583 μg/mL catalase, 42 μg/mL glucose oxidase, 1.7% w/v glucose, 1 mM DTT, 2 mM methyl viologen, 2 mM ascorbic acid, 1 μM Taxol, and 5 mM ATP) containing 0.5-3.0 nM kinesin motors (added in bulk). The coverslip was then mounted onto an indented microscope slide with imaging buffer and sealed using Twinsil two-component dental glue (Picodent). After a 5 min incubation at 37 °C to allow for hardening of the dental glue, the sample was brought to the microscope for imaging.

### MOTOR-PAINT IMAGING

Imaging was performed immediately after sample preparation at room temperature (20-23 °C) on a Nikon Ti-E inverted microscope equipped with a 100 x Apo TIRF oil immersion objective (NA. 1.49), a Perfect Focus System 3 (Nikon), and custom optics allowing for a tunable angle of incidence to perform (pseudo-) total internal reflection fluorescence (TIRF) microscopy. A Lighthub-6 laser combiner (Omicron) with a 638 nm laser (BrixX 500 mW multi-mode, Omicron) and a 405 nm laser (LuxX 60 mW, Omicron) was used for excitation. Emission light was separated from excitation light using a quad-band polychroic mirror (ZT405/488/561/640rpc, Chroma) and a quad-band emission filter (ZET405/488/561/640 m, Chroma). Detection was done using a Hamamatsu Flash 4.0v2 sCMOS camera. Image stacks of motors were acquired with a 60 ms exposure time, 12-13% laser power, and 15000–20000 images per field of view. Components were controlled using µManager (Edelstein et al., 2014).

### IDENTIFICATION OF CELL STAGE

Cells were classified as belonging to different stages based on their morphology as determined from the phalloidin staining. The number of neurites, their (relative) lengths, how branched they were, and the presence/absence of a remaining lamellipodium were used to assign neurons into the different stages. Stage 1 neurons had no neurites and a large lamellipodium. Stage 2a neurons most commonly had 1 to 4 neurites, all < 20 μm in length with no branches, and sometimes had a remaining lamellipodium. Stage 2b neurons typically had more and slightly longer neurites (up to ∼50 μm in length) with one or two branches, with no single neurite significantly longer than the rest. Stage 3 neurons had one (or sometimes two in our culture system) neurite(s) much longer than the rest that often had a few branches along it. Stage 4 neurons had extensive dendritic branching in addition to the presence of an axon.

### MOTOR-PAINT ANALYSIS

Preceding motor localization, raw movies were processed by median filtering to remove static and very rapidly moving (i.e. diffusive) particles (https://github.com/HohlbeinLab/FTM2). Subsequently, motors were detected and tracked using the Laplacian of Gaussian (LoG) detector and overlap tracker from TrackMate (Tinevez et al., 2017). The output data tables were converted into Results tables compatible with DoM for further processing.

For drift correction, 2 or 3 Tetraspeck beads were chosen per field of view and localized using Detection of Molecules (DoM) version 1.2.5 (https://github.com/UU-cellbiology/DoM_Utrecht). These localizations were then filtered and processed as follows. In the case of multiple bead localizations in the same frame, only the one with the highest signal-to-noise ratio (SNR) was kept. Outliers (>3 standard deviations away from the mean of the localizations in adjacent frames) were removed and replaced by a copy of the preceding localization. Subsequently, jumps of >150 nm were detected which represent abnormal drift. A rolling window was then applied between jumps to remove fluctuations. After correcting for jumps, the average of the first 1000 bead localizations was subtracted from all the localizations to create a drift correction table that was then applied to the motor localizations.

Motor tracks were then filtered. Tracks were only kept if they had ≥ 6 localizations, covered a distance > 200 nm, had a speed of 100-1500 nm/s and a ratio of the net:total displacement of 0.1-0.9. Subsequently, these tracks were filtered for angle to remove pauses or reversals in orientation (i.e. due to erroneously linked spots). This was done by calculating displacement vectors from every localization i to localization i+3. The angle between each pair of consecutive vectors was then calculated and stretches of ≥ 4 frames in which this angle was < 50° were retained.

The resulting particle table was then split into 4 separate tables depending on whether the track had a positive or negative displacement in x and y and each orientation was assigned a lookup table: tracks moving to the top-left were cyan, tracks to the top-right were blue, tracks to the bottom-right were magenta, and tracks to the bottom-left were yellow. Each of these particle tables were reconstructed separately using DoM and merged.

Alternatively, for the neurite-based colour-coding, further analysis was performed. A lasso selector tool was written in Python to select all particle localizations in a given primary neurite or its branches. Each neurite/branch was then rotated approximately horizontal, and the localizations from all the tracks within this neurite/branch were treated as a whole, median filtered and spline fit to produce a line used as the midline of that neurite/branch. After determining which end of this midline is closer to the soma, the localizations of each track were projected onto the midline and the start- and end-point of the track were compared to determine whether the track was moving towards or away from the soma along the midline. Subsequently, tracks from the branches of a given neurite were grouped to report a single value per neurite (including all its branches). Minus-end-out fractions were calculated as the number of localizations in minus-end-out tracks over the total number of localizations in that neurite. Values from filopodia (very thin protrusions < 10 µm in length) were excluded, as well as from axons and axon-like neurites (> 90% plus-end-out) where indicated. These data were then exported and plotted using GraphPad Prism version 10.1.1.

Codes are available at https://github.com/UU-cellbiology/neuron_motorPAINT.

### ULTRASTRUCTURE EXPANSION MICROSCOPY (U-EXM)

Neurons were fixed differently depending on whether the centrioles or neurites were being imaged. For imaging neurites, soluble tubulin was extracted by incubating cultured neurons for 1 minute with 0.3% v/v TritonX-100, 1 M sucrose, and 0.1% w/v glutaraldehyde in MRB80 and then fixed for 8 min with 2% w/v paraformaldehyde and 1 M sucrose in MRB80. Cells were then quenched twice for 5 min with 100 mM sodium borohydride in PBS and washed with PBS prior to proceeding. For imaging of the centrioles, neurons were incubated with pre-warmed MRB80 for 1 minute, followed by fixation in ice-cold methanol and 6 mM EGTA on ice for 12 minutes. Hereafter, neurons were rehydrated using four PBS washes (60 seconds, 2 minutes, 5 minutes, 5 minutes).

Anchoring was performed by incubating fixed neurons in 1.4% w/v formaldehyde and 3% w/v acrylamide (Sigma-Aldrich, A4058) in PBS for 3 hours at 37 °C. After several PBS washes, neurons were incubated with the U-ExM gelation solution (19% w/v sodium acrylate, 10% w/v acrylamide (Sigma-Aldrich, A4058), 0.1% w/v N,N′-methylenebisacrylamide (Sigma-Aldrich M1533), 0.5% APS w/v (Sigma-Aldrich, 215589), 0.5% w/v TEMED (Bio-Rad, 1610800), and 1x PBS) for 2-3 minutes on ice and then polymerized for 1 hour at 37 °C (Gambarotto et al., 2019). The 38% w/v sodium acrylate stocks were prepared as previously described (Damstra et al., 2022). Briefly, acrylic acid (Sigma-Aldrich, 147230) was neutralized with 10 M sodium hydroxide to a final pH of 7.5– 8 and stored at −20 °C. Gels were denatured in the presence of 345 mM SDS, 225 mM Tris (pH 9), and 500 mM NaCl for 1.5 hours at 95 °C. After denaturation, gels were washed two times for at least 30 minutes in ultrapure water, followed by two washes in PBS for at least 30 minutes with shaking. The expansion factor used for image calibration was calculated by dividing the gel diameter after several MilliQ washes (until the gels did not expand any further) by the coverslip size. For post-expansion labelling, gels were then cut into pieces and incubated with primary antibodies (see antibody table) diluted 1:250 in PBS supplemented with 2% w/v BSA and 0.1% w/v TritonX-100 overnight at room temperature with shaking. Hereafter, gels were washed 4x 15 minutes in PBS supplemented with 0.1% v/v TritonX-100. Gels were then incubated with secondary antibodies (see antibody table) diluted to 1:500 in PBS supplemented with 2% w/v BSA and 0.1% v/v TritonX-100 for 3 hours at room temperature with shaking and finally washed 4x 15 minutes in 0.1% v/v TritonX-100 in PBS.

Gels were mounted on 25 mm, No. 1.5H poly-L-lysine coated coverslips in an imaging ring and imaged on a Leica TCS SP8 STED 3X microscope using a 63x/1.2 NA water-immersion objective. A white laser, pulsed at 80 MHz, was used for excitation. Leica GaAsP HyD hybrid detectors were used for fluorescence detection. Using Las X software, stacks with a lateral pixel size of 70-75 nm and an axial spacing of 180 nm were acquired using the frame-sequential mode.

### OTHER ANALYSIS AND IMAGE PROCESSING

Confocal, TIRF, and motor-PAINT images were processed in FIJI (ImageJ) version 1.54f. In all instances, brightness and contrast were adjusted linearly. The background of the kymograph in Figure 4D was subtracted using a rolling ball radius of 20 and no smoothing. Images of neurites were made from the whole field of view using Edit > Selection > Straighten separately for each channel. Line profiles were created using Analyze > Plot profile for each channel using a line width of 10 pixels (∼200 nm) and exporting the resulting list of values for plotting in GraphPad Prism version 10.1.1. Note that for line plots, data were normalized separately for each channel using the minimum and maximum values in that dataset.

Expansion images were drift corrected and deconvolved with Huygens Professional (v17.04, Scientific Volume Imaging), using the classic maximum likelihood estimation (CMLE) algorithm. Images were iterated with CMLE over 8-10 iterations with a signal-to-noise ratio of 15. In all cases except for the whole cell image of the stage 2a neuron shown in Figure 3C, maximum projections were made using approximately 3-5 slices around the middle of the neurites or approximately 6-9 slices around the centrioles. For the whole cell image of the stage 2a neuron in Figure 3C, a sum projection was used instead.

To trace individual microtubules emanating from centrioles in expanded neurons, deconvolved expansion image z-stacks were cropped to a quadratic area with sides of 8 µm (1.78 µm when corrected for expansion factor) centred around the centrioles in ImageJ (v1.54f). Cropped stacks were loaded into BigTrace (v0.39, https://github.com/ekatrukha/BigTrace) to verify their attachment to the centrioles in 3D. Microtubules that were clearly attached to the centrioles were traced manually in the 2D max projections for easy visualization.

To analyze the radial distribution of acetylated and tyrosinated microtubules in expanded neurites, deconvolved image stacks were processed using custom scripts in ImageJ (v1.54f) and MATLAB (R2024b) as described in detail elsewhere (Katrukha et al., 2021). Briefly, on maximum intensity projections (XY plane), we drew polylines of sufficient thickness (300 px) to segment out neurite portions 44 µm (10 µm when corrected for expansion factor) in length proximal to the cell soma. Using Selection > Straighten on the corresponding z-stacks generated straightened B-spline interpolated stacks of the neurite sections. These z-stacks were then resliced perpendicularly to the neurite axis (YZ-plane) to visualize the neurite cross-section. From this, we could semi-automatically find the boundary of the neurite in each slice using first a bounding rectangle that encompasses the neurite (per slice) and then a smooth closed spline (approximately oval). To build a radial intensity distribution from neurite border to center, closed spline contours were then shrunken pixel by pixel in each YZ-slice while measuring ROI area and integrated fluorescence intensity. From this, we could ascertain the average fluorescence intensity per contour iteration, allowing us to calculate a radial intensity distribution by calculating the radius corresponding to each area (assuming the neurite cross-section is circular).

## ACKNOWLEDGEMENTS

This work was supported by the European Research Council (Consolidator Grant 819219-CellularLogistics to L.C.K) and by the Dutch Foundation for Scientific Research (NOW-VICI Grant VI.C.212.062 to L.C.K).

## AUTHOR CONTRIBUTIONS

M.K.I. and L.C.K. designed the study. M.K.I. performed motor-PAINT experiments and developed and performed analysis, as well as performed live-cell experiments and analysis. A.K.S. performed expansion microscopy experiments. H.N.V. performed motor-PAINT experiments and developed and performed analysis. B.C.D. performed motor-PAINT experiments and analysis. M.K.I. and L.C.K. wrote the manuscript with input from all other authors. L.C.K. supervised the study.

## AUTHOR NOTES

### Disclosures

The authors declare no competing interests exist.

---

**Figure S1:**
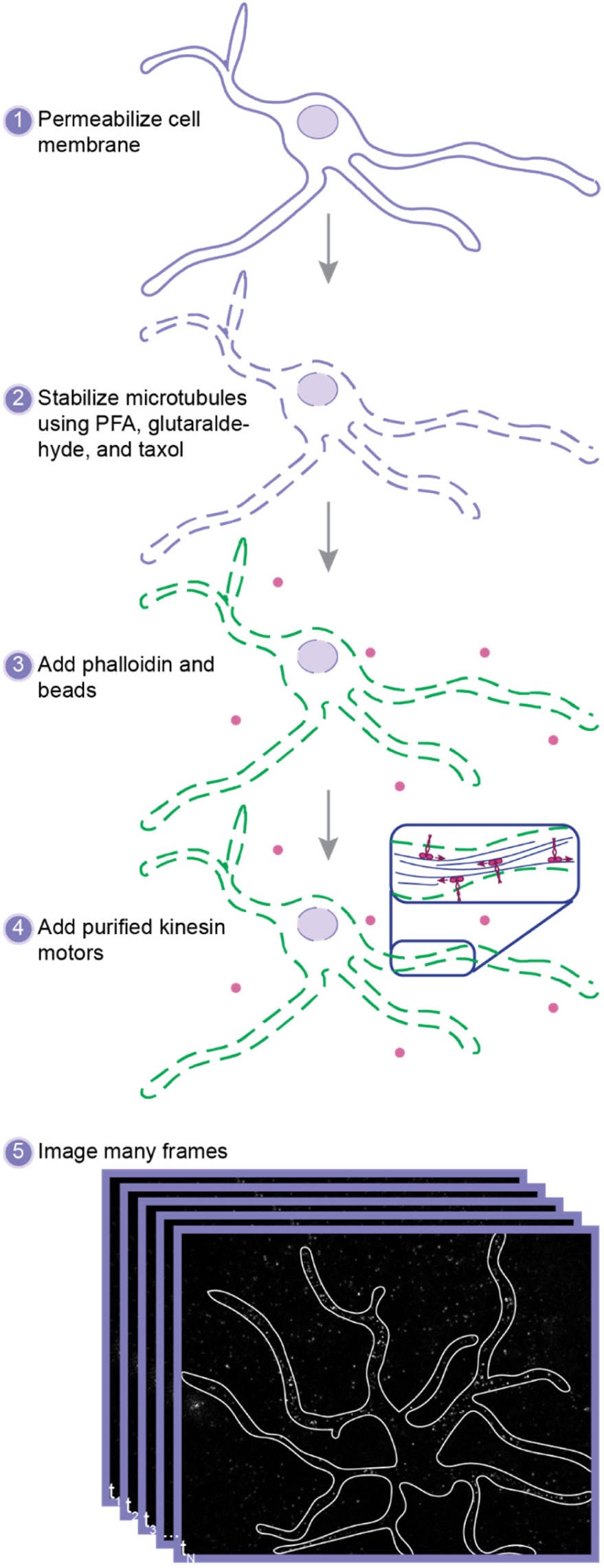
Schematic showing the motor-PAINT workflow. Neurons are briefly permeabilized (1) and then exposed to low concentrations of PFA and glutaraldehyde for a gentle fixation (2). After washing with a Taxol-containing buffer, neurons are briefly incubated with phalloidin and multicolour beads (3). Subsequently, purified truncated kinesin motors labeled with JF646 are added in an imaging buffer (4), the chamber is sealed, and each cell is imaged for many thousands of frames (5) to allow super-resolutions reconstructions of the microtubules and their orientations.

**Figure S2:**
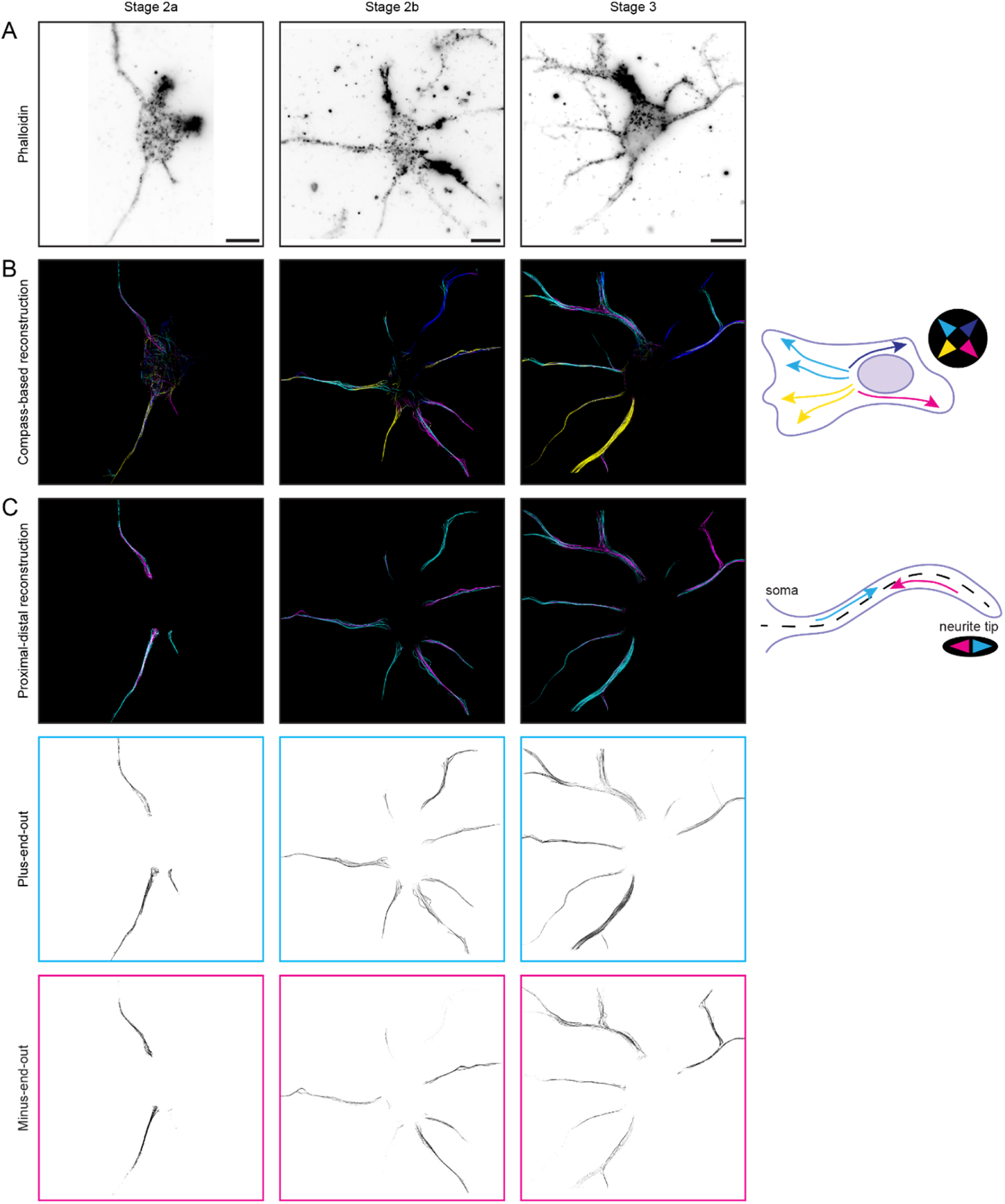
Different views of the same cells shown in Figure 1C. **A)** Phalloidin stainings of these neurons showing neuron morphology as used to classify cells into stage 2a, 2b or 3. Scale bars 10 µm. **B)** The total tracks after filtering showing the microtubules in these neurons, but colour-coded based on whether the tracks move towards the top-left, (cyan), top-right (blue), bottom-right (magenta) or bottom-left (yellow). **C)** The tracks in each of the neurites of these neurons colour-coded based on whether they are moving towards (magenta) or away from (cyan) the soma. Single-channel images are shown below.

**Figure S3:**
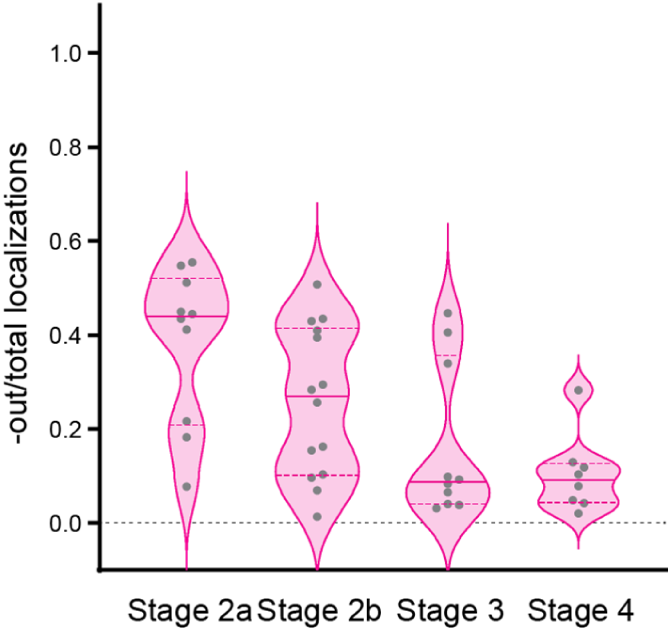
Summary of the data for cells in which the longest neurite could be clearly identified. Quantification of the fraction of localizations constituting minus-end-out tracks over the total amount of localizations for the longest neurites/axons from neurons in stages 2a, 2b, 3, and 4. Each dot represents one neurite. Medians (0.44, 0.27, 0.09, 0.09) and interquartile ranges ((0.21, 0.52), (0.10, 0.41), (0.04, 0.36), (0.04, 0.13)) are shown. n = 10, 14, 10, 8 neurites from N = 10, 14, 10, 8 cells for stages 2a, 2b, 3, 4.

**Figure S4:**
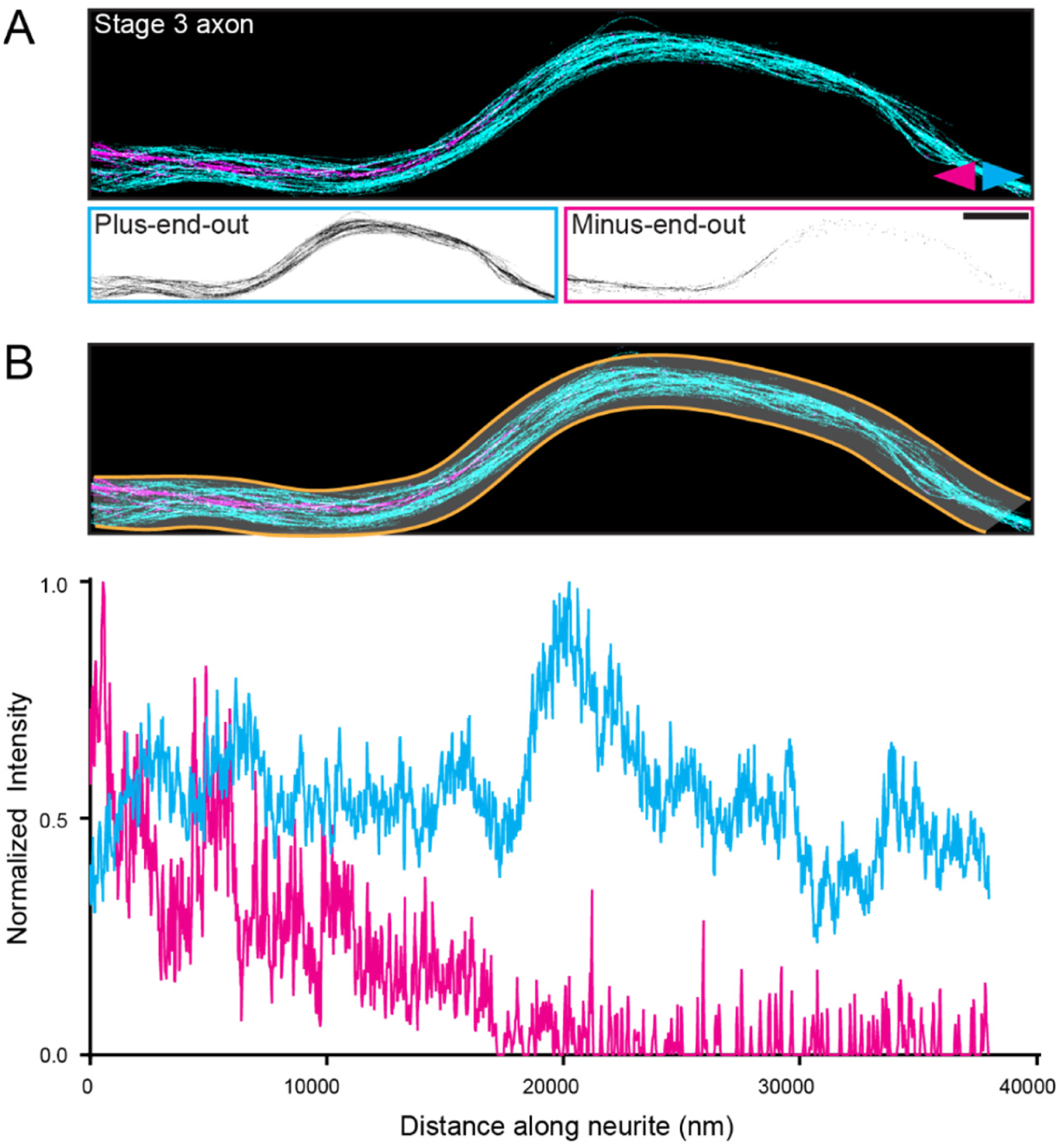
Some axon-like neurites and axons in stage 2b and stage 3 cells retain a proximal bundle of minus-end-out microtubules. **A)** A motor-PAINT reconstruction of an axon/axon-like neurite from a stage 3 cell with microtubules pseudo-coloured based on whether their plus-end is oriented towards (magenta) or away from (cyan) the soma. Single channel images are shown below. Scale bars 2 µm. **B)** Intensity profiles of minus-end-out (magenta) and plus-end-out (cyan) microtubules along a line thick enough to enclose the entire neurite (between the yellow lines shown along the neurite). Normalization was done independently for the two orientations using the minimum and maximum values of those data sets.

**Figure S5:**
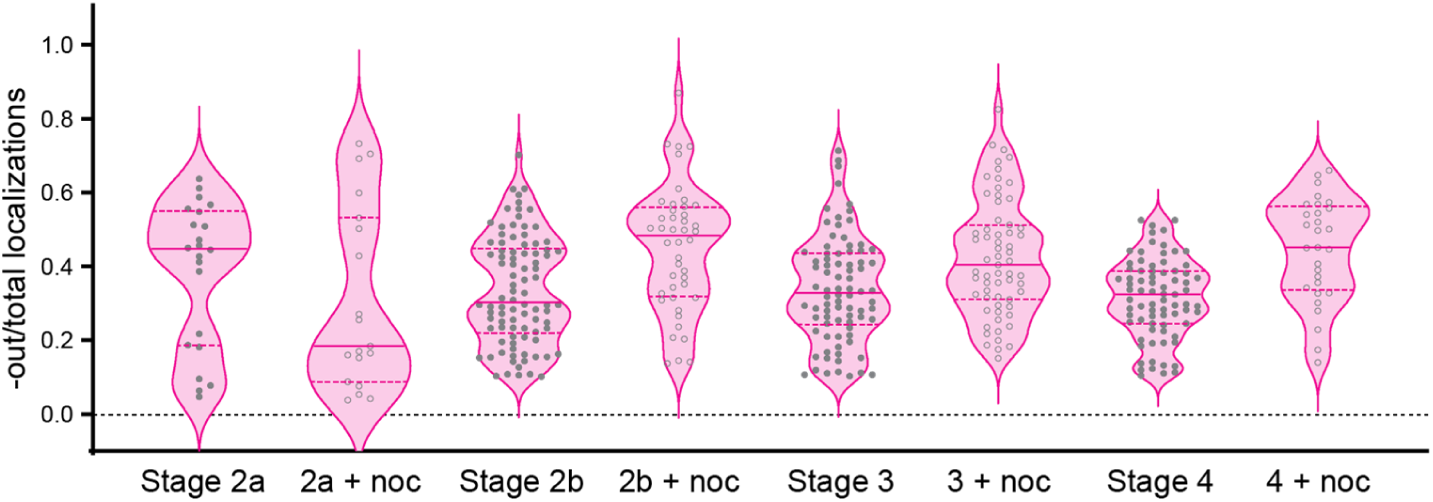
Summary of the data throughout excluding axons and axon-like neurites. Quantification of the fraction of localizations constituting minus-end-out tracks over the total amount of localizations for neurons in stages 2a, 2b, 3, and 4 without and with nocodazole. Each dot represents one neurite. Medians (0.45, 0.18, 0.30, 0.48, 0.33, 0.40, 0.32, 0.45) and interquartile ranges ((0.19, 0.55), (0.09, 0.53), (0.22, 0.45), (0.32, 0.56), (0.24, 0.44), (0.31, 0.51), (0.25,0.39), (0.34, 0.56)) are shown. n = 22, 19, 90, 44, 82, 63, 77, 29 neurites from N = 9, 8, 18, 11, 17, 14, 14, 6 cells for stages 2a without/with, 2b without/with, 3 without/with, and 4 without/with nocodazole.

**Figure S6:**
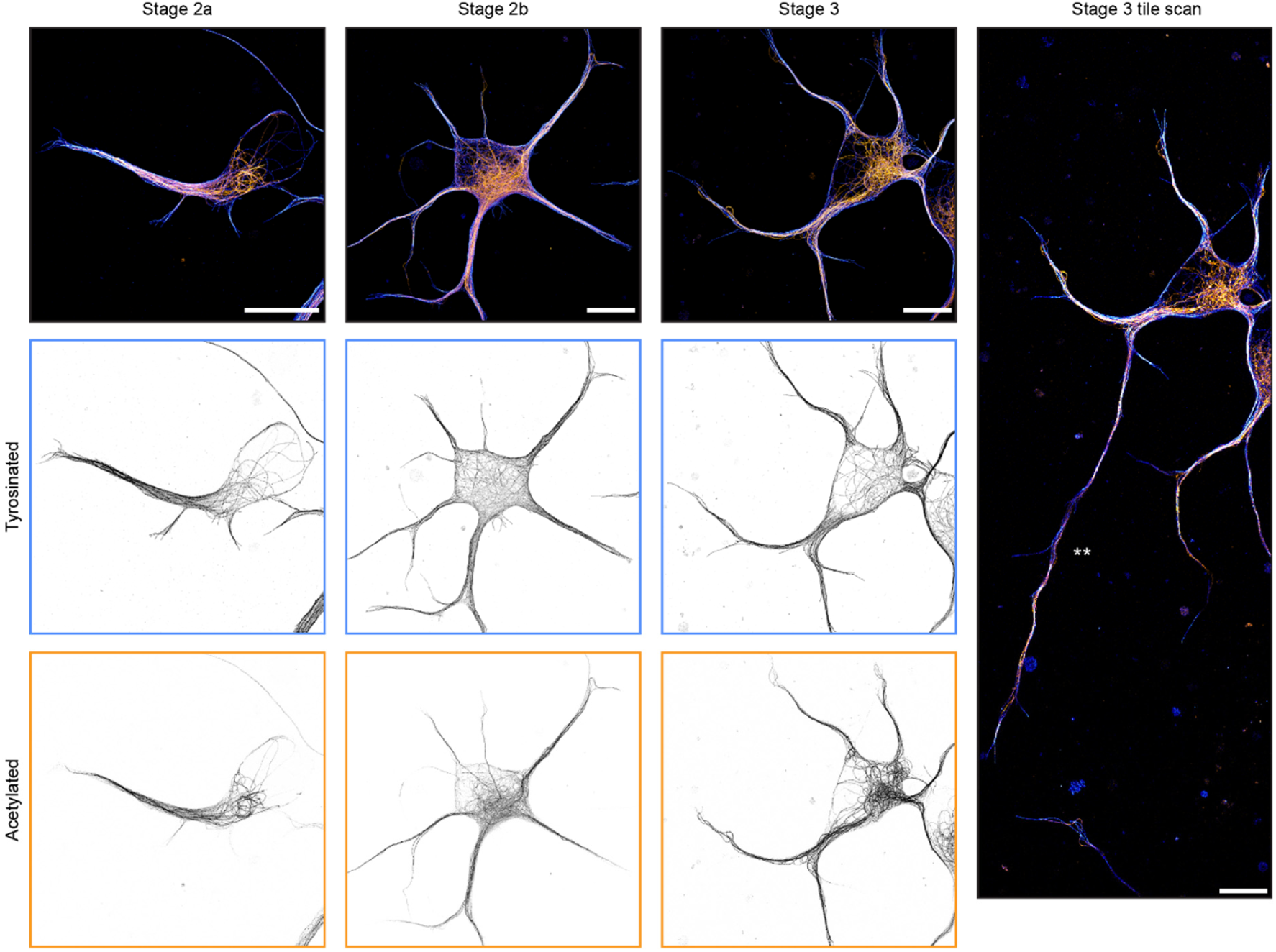
U-ExM images of the stage 2a, stage 2b, and stage 3 (left to right) neurons from Figures 2A, C, and E showing both acetylated (orange) and tyrosinated (blue) microtubules. Single channel images are shown below. Right: Tile scan of the stage 3 neuron. ** indicates developing axon. Scale bars 10 µm (corrected for expansion).

**Figure S7:**
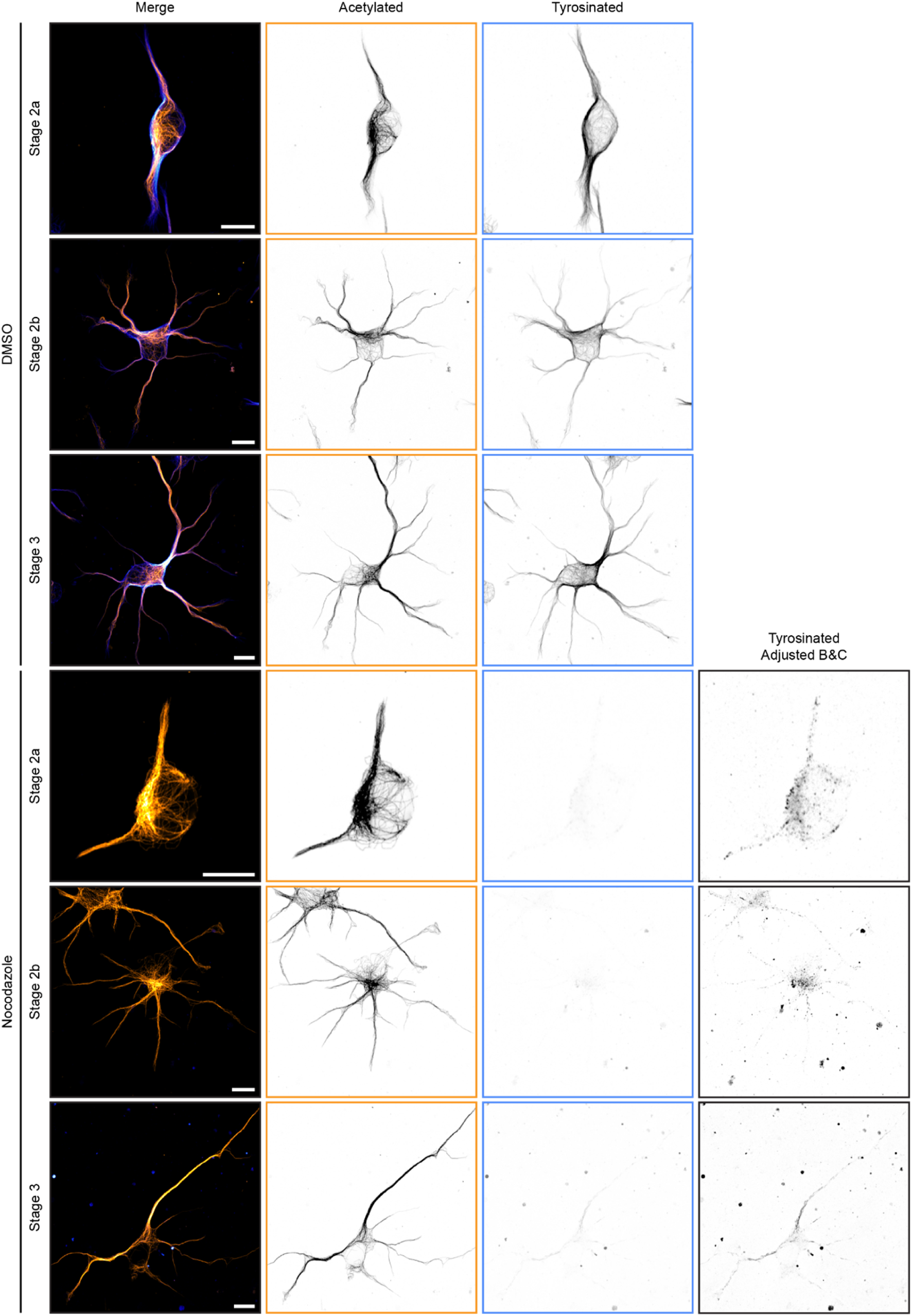
Nocodazole treatment is efficient and non-lethal in developing neurons. Examples confocal images showing tyrosinated (blue) and acetylated (orange) microtubules in stage 2a, 2b, and 3 neurons treated with 4 µM nocodazole for 2.5 hours or an equivalent amount of DMSO showing the reduction in intensity of tyrosinated microtubules after nocodazole treatment. Imaging conditions were kept the same in all conditions and the brightness and contrast were scaled the same except for in the ‘Adjusted B&C’ column, for which the brightness and contrast were greatly increased to show any remaining tyrosinated microtubule signal. Scale bars 10 µm.

**Figure S8:**
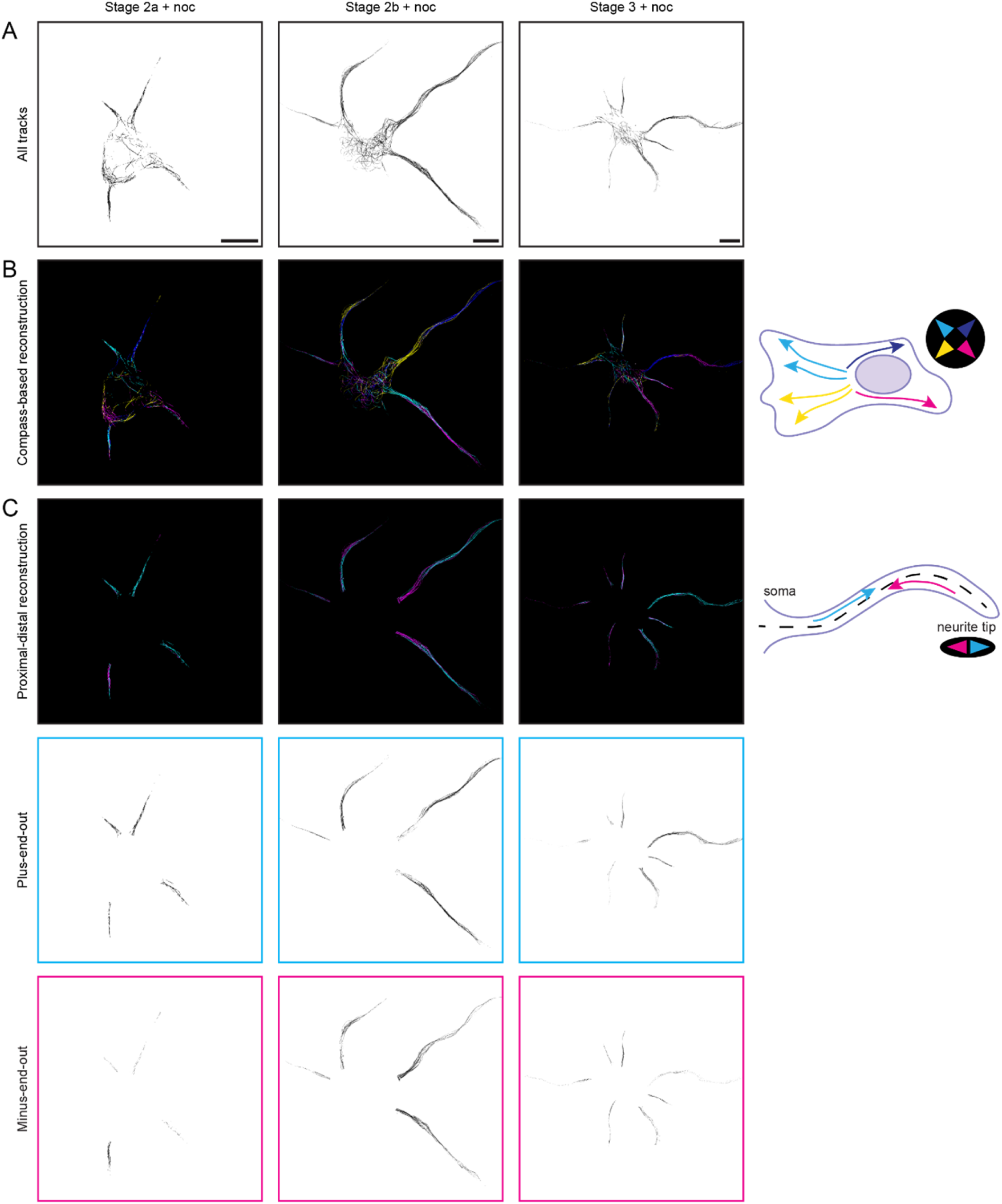
Different views of the same cells shown in Figures 2G, I and 3A. **A)** Total tracks after filtering (all orientations) showing all microtubules in these neurons. Scale bars 10 µm. **B)** The total tracks after filtering showing the microtubules in these neurons, but colour-coded based on whether the tracks move towards the top-left, (cyan), top-right (blue), bottom-right (magenta) or bottom-left (yellow). **C)** The tracks in each of the neurites of these neurons colour-coded based on whether they are moving towards (magenta) or away from (cyan) the soma. Single-channel images are shown below.

**Figure S9:**
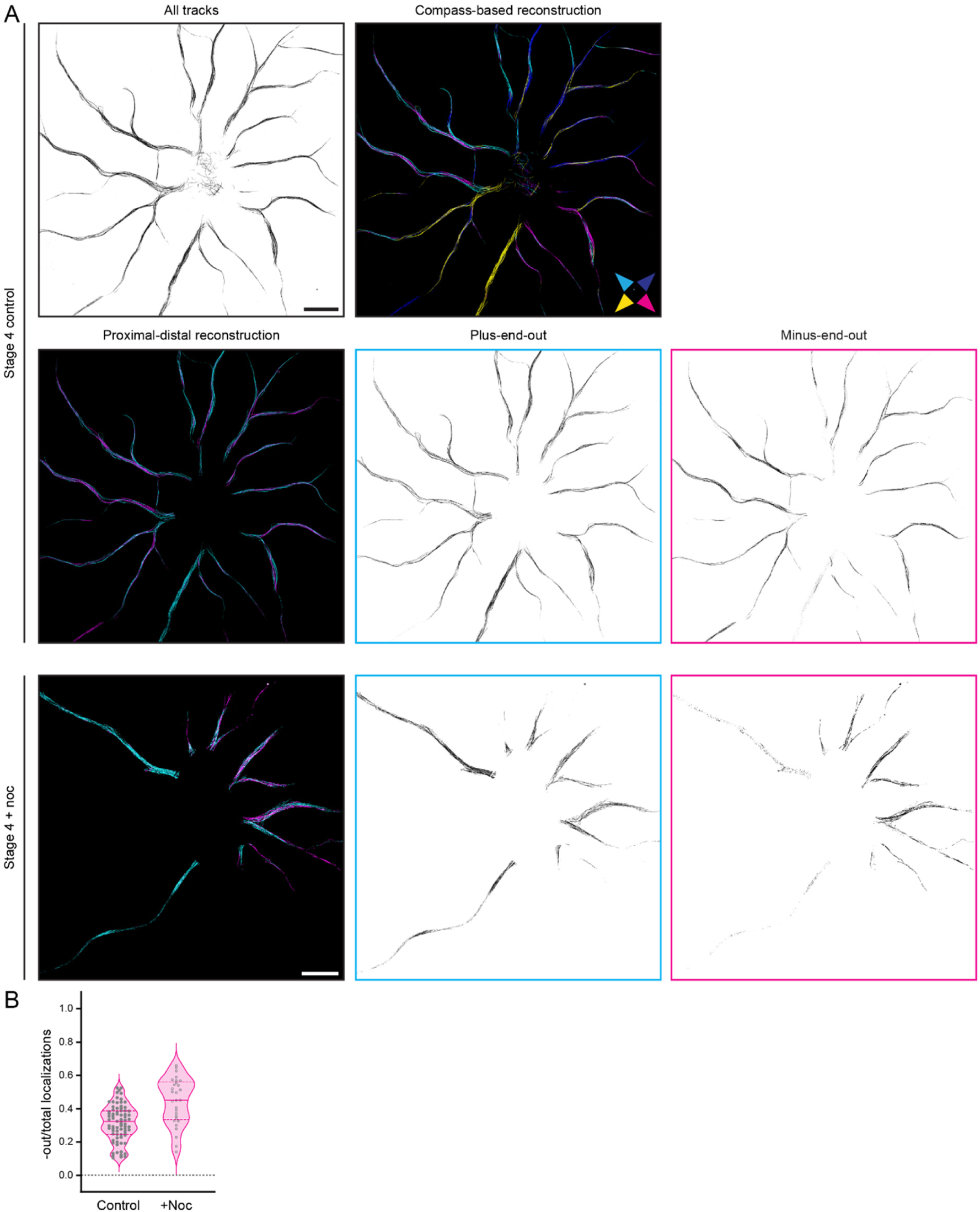
**A)** motor-PAINT reconstructions of a control stage 4 neuron and a stage 4 neurons after nocodazole treatment. Top: For the control neurons, the total tracks after filtering are shown, as well as these tracks colour-coded based on whether the tracks move towards the top-left, (cyan), top-right (blue), bottom-right (magenta) or bottom-left (yellow). Second row: The tracks in each of the neurites is shown colour-coded based on whether they are moving towards (magenta) or away from (cyan) the soma. Single-channel images are shown to the right. Bottom: For the nocodazole-treated neuron, the tracks in each of the neurites is shown colour-coded based on whether they are moving towards (magenta) or away from (cyan) the soma. Single-channel images are shown to the right. Scale bars 10 µm. **B)** Quantification of the fraction of localizations constituting minus-end-out tracks over the total amount of localizations for stage 4 neurons without (control) and with (+ noc) a nocodazole treatment. Each dot represents one neurite. Medians (0.28, 0.45) and interquartile ranges ((0.13, 0.37), (0.31, 0.55)) are shown. n = 94, 32 neurites from N = 14, 6 cells for control and nocodazole-treated cells.

**Figure S10:**
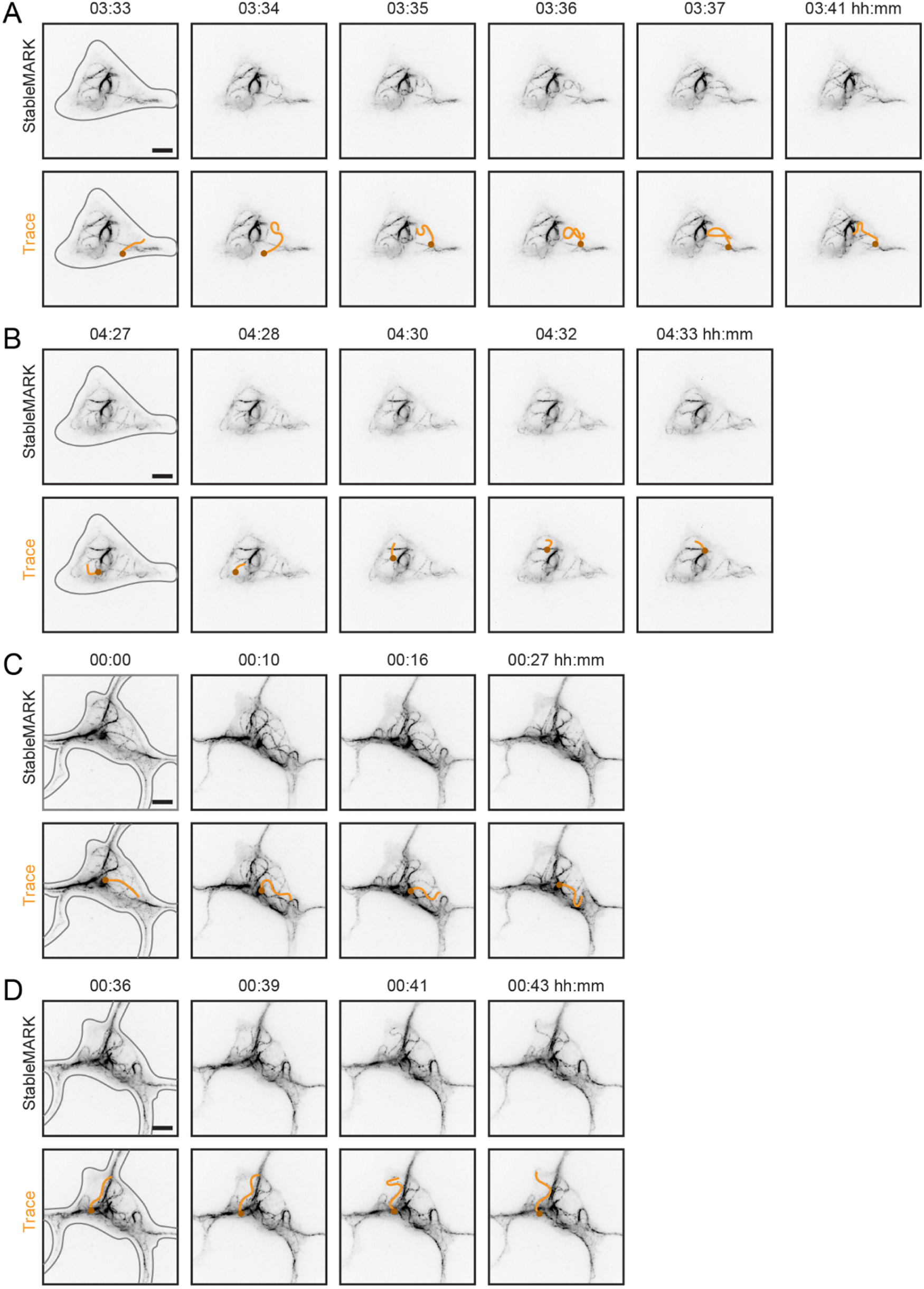
Live cell examples showing the highly dynamic stable microtubules in stage 2a (**A**, **B**) and stage 2b (**C**, **D**) neurons. Neurons were electroporated with StableMARK. Example microtubules are traced in orange in each of the bottom rows, with the same end marked by a small circle in each frame to indicate orientation. First frames also have the cell outline in grey. Times shown above each frame are in hours:minutes. Scale bars 5 µm. A and B correspond to Movie 2. C and D correspond to Movie 3.

## SUPPLEMENTARY MOVIES

**Movie 1:** An example of a stage 2b neuron expressing StableMARK showing the retrograde flow of stable microtubules in the neurites. Timestamps shown are in hours:minutes. Movie played at 10 fps. Scale bar 5 µm.

**Movie 2:** An example of a stage 2a neuron expressing StableMARK showing the very motile and sliding stable microtubules in this stage. Timestamps shown are in hours:minutes. Movie played at 10 fps. Scale bar 5 µm.

**Movie 3:** An example of a stage 2b neuron expressing StableMARK showing the very motile and sliding stable microtubules in this stage, as well as the clear link of a bundle of stable microtubules to a focal point (potentially the centrosome). Also shows curling back of stable microtubules in neurites to reverse orientation. Timestamps shown are in hours:minutes. Movie played at 10 fps. Scale bar 5 µm.

**Movie 4:** An example of a stage 2b neuron expressing StableMARK showing the very stable link of a bundle of stable microtubules to the centrosome. Timestamps shown are in hours:minutes. Movie played at 10 fps. Scale bar 5 µm.

**Movie 5:** An example of a stage 2b neuron expressing StableMARK showing two means by which stable microtubules can reverse their orientation: sliding from one neurite into another and reversing within the same neurite. Timestamps shown are in hours:minutes. Movie played at 10 fps. Scale bar 5 µm.

## Notes

### Competing Interest Statement

The authors have declared no competing interest.

